# Conformational changes in mitochondrial complex I from the thermophilic eukaryote *Chaetomium thermophilum*

**DOI:** 10.1101/2022.05.13.491814

**Authors:** Eike Laube, Jakob Meier-Credo, Julian D. Langer, Werner Kühlbrandt

## Abstract

Mitochondrial complex I is a redox-driven proton pump that generates proton-motive force across the inner mitochondrial membrane, powering oxidative phosphorylation and ATP synthesis in eukaryotes. We report the structure of complex I from the thermophilic fungus *Chaetomium thermophilum*, determined by cryoEM up to 2.4 Å resolution. We show that the complex undergoes a transition between two conformations, which we refer to as form 1 and 2. The conformational switch is manifest in a twisting movement of the peripheral arm relative to the membrane arm, but most notably in substantial rearrangements of the Q-binding cavity and the E-channel, resulting in a continuous aqueous passage from the E-channel to subunit ND5 at the far end of the membrane arm. The conformational changes in the complex interior resemble those reported for mammalian complex I, suggesting a highly conserved, universal mechanism of coupling electron transport to proton pumping.

## Introduction

Mitochondrial NADH:ubiquinone oxidoreductase (complex I) is the major entry point of electrons into the respiratory chain *(1, 2)*. Complex I is a multi-subunit membrane protein complex consisting of around 45 different protein subunits. It couples the transfer of two electrons from reduced nicotinamide adenine dinucleotide (NADH) to ubiquinone (Q) to the transfer of four protons across the inner mitochondrial membrane *(3-5)*. Electron and proton transfer accounts for a large part of the proton-motive force that drives ATP synthesis by the F-ATP synthase in the same membrane *(6)*.

A minimal version of mitochondrial complex I is found in bacteria. The bacterial complex consists of 14 strictly conserved core subunits that are sufficient for catalytic activity. In the course of eukaryotic evolution, ∼30 accessory subunits have been incorporated into complex I, increasing its molecular mass from ∼550 kDa to ∼1 MDa. Although the exact roles of most accessory subunits are unknown, some have been shown to regulate complex I assembly, stability and activity *(7)*.

Mitochondrial complex I has a characteristic L-shape, with a membrane arm embedded in the lipid bilayer of the inner mitochondrial membrane, joined at one end to the peripheral arm that extends into the mitochondrial matrix. The peripheral arm transfers electrons from NADH to Q, via a bound flavin mononucleotide (FMN) along a chain of eight iron-sulphur (Fe-S) clusters *(8-11)*. The membrane arm pumps protons across the inner mitochondrial membrane. It contains three antiporter-like subunits that form an elongated hydrophilic passage with numerous internal water molecules. Of these, only the antiporter-like subunit ND5 at the distal end of the membrane arm appears to have an aqueous connection to the intermembrane space (IMS) *(12, 13)*. The Q substrate binds in a cavity above the level of the membrane surface near the junction of the two arms, where it is reduced to ubiquinol (QH_2_) by Fe-S cluster N2 *(14)*. An aqueous tunnel referred to as the E-channel is lined by strictly conserved glutamate residues of core subunits ND1, ND3 and ND4L. The E-channel connects the Q-binding cavity to the internal hydrophilic passage within the membrane arm.

Recent electron cryo-microscopy (cryoEM) structures of complex I show that the Q headgroup binds at two primary sites within the cavity *(13, 15, 16)*. The deeper binding site (referred to as Q_d_) is located next to N2. The Q_s_ site is located in the shallower part of the cavity close to its entrance. Presumably, binding of Q, QH_2_ or Q intermediates at different binding sites within the Q-binding cavity during the catalytic cycle is linked to proton translocation *(12, 13, 15)*, but the exact molecular mechanism is not understood.

Mammalian mitochondrial complex I incubated at elevated temperatures (>30 °C) without NADH and Q substrates undergoes a reversible transition from an active to a deactive state, characterized by a significant decrease in turnover rate *(17, 18)*. The deactive state is thought to protect the complex against the production of reactive oxygen species (ROS) generated by reverse electron transfer during reperfusion in ischemic tissues *(19, 20)*. When fresh substrate is added, complex I reverts to the active state. The transition between the active and deactive state is characterized by a lag phase in NADH:Q oxidoreductase activity that lasts up to 2 min *(21)*, before NADH is again oxidized at a linear rate *(17)*.

CryoEM of mammalian *(13, 15, 22, 23)* and plant complex I *(24)* has resolved two different main conformations, characterized by a tilt of the peripheral arm relative to the membrane arm. In the mammalian complex, internal rearrangements have been described in the Q-binding cavity and at transmembrane helix (TMH) 3 of core subunit ND6 *(13, 22)*. At present it is not clear how these two conformations or the associated internal rearrangements relate to the mechanism of electron transport and proton translocation, whether they represent the deactive and active state of complex I *(22)*, or are substates of the active state and thus represent different catalytic intermediates *(13)*.

In the present study we investigate mitochondrial complex I from the filamentous fungus *Chaetomium thermophilum*, one of the few well-characterized thermophilic eukaryotes. *C. thermophilum* has an optimal growth temperature of 50-55 °C, but tolerates temperatures up to 60 °C *(25)*. It grows naturally in dung or compost heaps, but is easily cultivated in standard media. Proteins and macromolecular complexes from thermophiles are good targets for structural biology, as they are usually more robust than their mesophilic counterparts *(26, 27)*, and hence more easily purified and crystallized. The properties of thermostable proteins, in particular their higher resistance to denaturation, are equally beneficial for cryoEM. We determined the cryoEM structure of mitochondrial complex I from *C. thermophilum* at 2.4 Å resolution. Even though we expected this thermotolerant complex to be more rigid than complex I from mammals or plants, we found that it undergoes major conformational changes. Structures of three different conformations were resolved, which we refer to as form 1 and form 2, plus an inhibited conformation.

## Results

### Subunit composition and overall structure of *C. thermophilum* complex I

Complex I from *C. thermophilum (Ct*-complex I) was solubilized either in the detergent *n*-dodecyl β-D-maltoside (DDM), or in lauryl maltose neopentyl glycol (LMNG). The 2.4 Å map allowed us to model the 14 core subunits and 29 accessory subunits with confidence (**Fig. 1**). For subunit assignment and atomic model building, we initially used the cryoEM structure of complex I from the aerobic yeast *Yarrowia lipolytica (Yl*-complex I; PDB 6RFR) *(16)*. For consistency, we adopted the subunit nomenclature of human complex I, except for the accessory subunit NUXM of the membrane arm, which is absent in mammals.

**Fig. 1.**
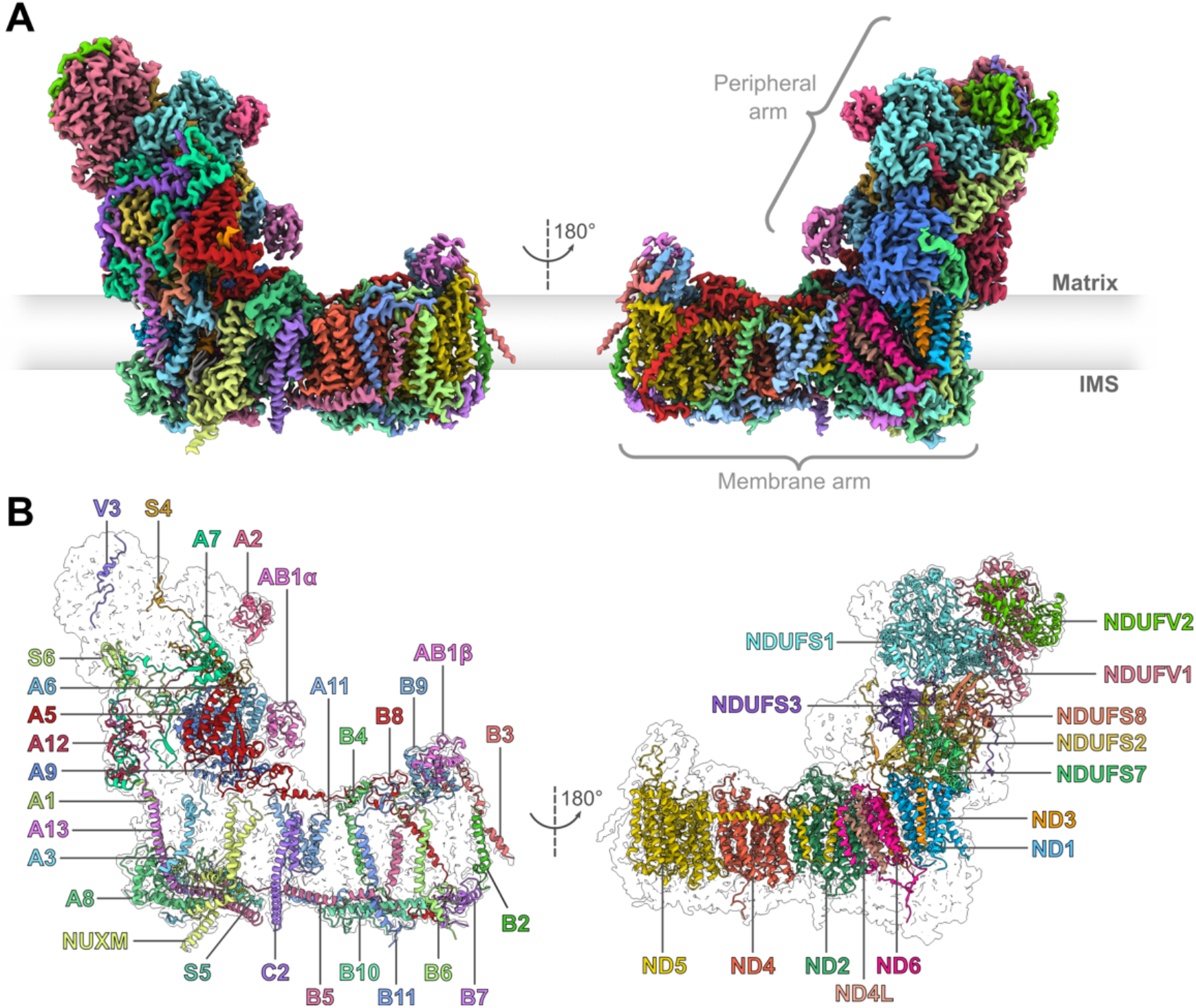
CryoEM density map and modelled structure of mitochondrial complex I from *C. thermophilum*. **(A)** CryoEM density map colored by subunit (IMS: Intermembrane space). **(B)** The 29 accessory (left) and 14 core (right) subunits of *Ct*-complex I. The subunit nomenclature is that of human complex I. The prefix ‘NDUF’ of accessory subunits was omitted for clarity. The subunit NUXM is not present in human complex I and named in accordance with complex I from *Yarrowia lipolytica*.

The overall structure of *Ct*-complex I is similar to that of *Y. lipolytica (28)*. The subunit composition is well-conserved, except for an accessory subunit at the top of the peripheral arm, binding at the same site as subunit NDUFV3, with which it shares weak sequence similarity. We assume that its role is analogous to that of mammalian NDUFV3 and refer it to as *Ct*-NDUFV3 (**Fig. S1**).

Subunits NDUFB2 and NDUFA1 were not annotated in the genome of *C. thermophilum (29)*. To determine their sequences as well as that of *Ct*-NDUFV3, we initially modelled these three subunits as polyalanine. Subsequently, map features indicating particular sidechains were used to place matching residues. The resulting peptide fragments served as query sequences for BLAST searches in a 6-frame-translation database from the *C. thermophilum* genome. Missing residues were then modelled according to the amino acid sequence of the returned hits. In this way all subunits of *Ct*-complex I were modelled and finally confirmed by MS.

Apart from the 43 protein subunits, the structure of *Ct-*complex I resolves thirteen bound cofactors (**Fig. 2A**). The peripheral arm contains eight Fe-S clusters (six Fe_4_S_4_ and two Fe_2_S_2_), a coordinated Zn^2+^ ion, one FMN, one nicotinamide adenine dinucleotide phosphate (NADPH) and a phosphopantetheine group, covalently attached to subunit NDUFAB1α. NDUFAB1 belongs to the group of mitochondrial acyl carrier proteins, which are involved in mitochondrial fatty acid synthesis *(30, 31)* and the assembly and activation of respiratory chain complexes *(32)*. *Yl*-complex I contains two isoforms of subunit NDUFAB1 (ACPM in *Y. lipolytica)*, one in the peripheral arm (ACPM1) and one at the tip of the membrane arm (ACPM2). In *C. thermophilum*, both positions are occupied by identical copies, referred to as NDUFAB1α and β.

**Fig. 2.**
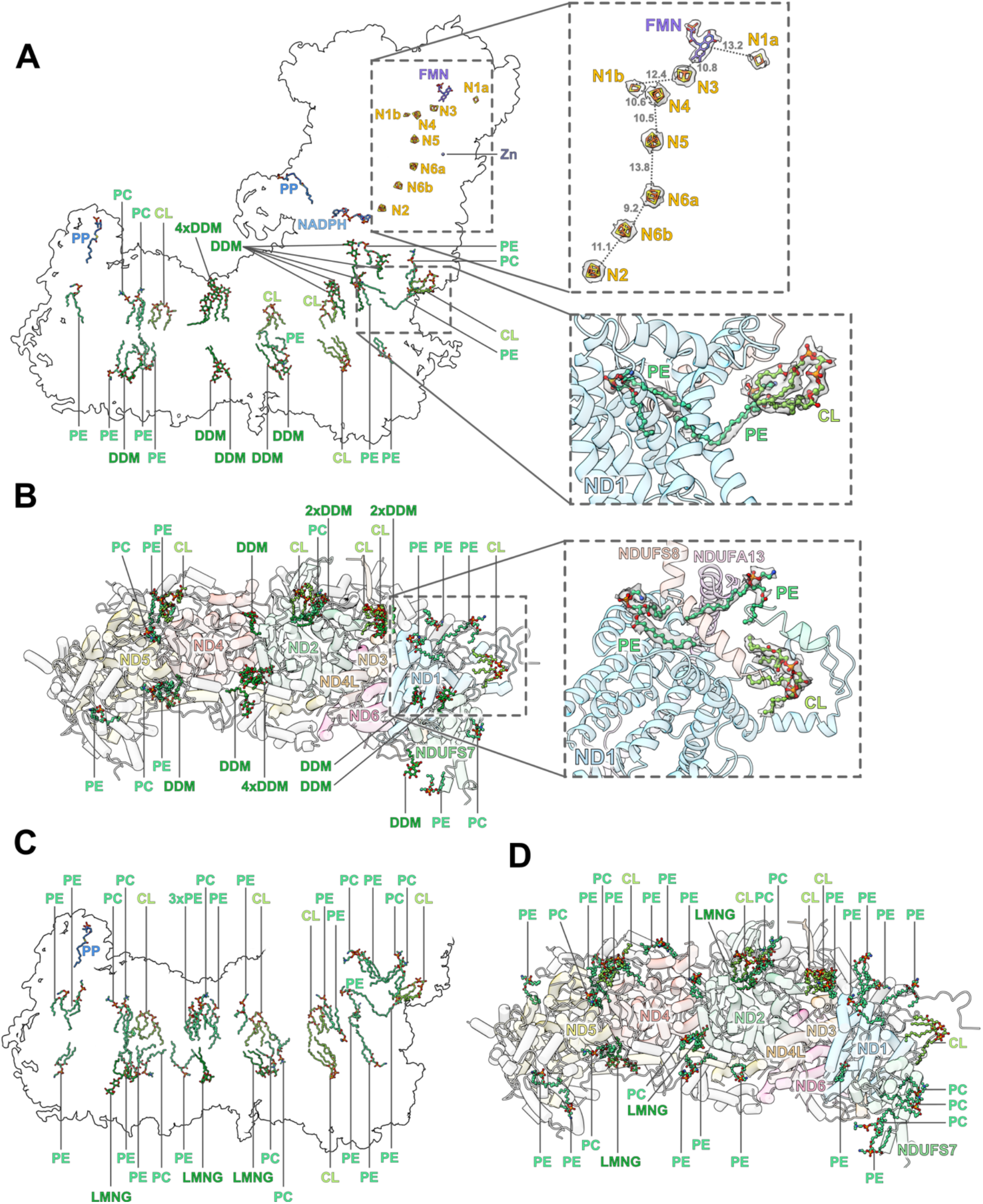
Cofactors and lipids in *C. thermophilum* complex I. **(A)** Cofactors, lipids and detergent molecules identified in the map of DDM-solubilized *Ct*-complex I. Upper inset: Distances (in Å) along the redox chain between FMN and the Fe-S clusters (N). Lower inset: Fatty-acid tails of lipids near subunit ND1 run almost parallel to the membrane plane. **(B)** View from the mitochondrial matrix side on the membrane arm of complex I in DDM (peripheral arm omitted for clarity). Inset: Top view of tilted lipids near subunit ND1. **(C)** Lipids and detergent molecules in the membrane arm of LMNG-solubilized *Ct*-complex I. **(D)** View of the membrane arm in LMNG from the matrix side (peripheral arm omitted for clarity). FMN: Flavin mononucleotide, NADPH: Nicotinamide adenine dinucleotide phosphate, CL: Cardiolipin, PE: Phosphatidylethanolamine, PC: Phosphatidylcholine, PP: Phosphopantetheine, DDM: n-dodecyl β-D-maltoside, LMNG: lauryl maltose neopentyl glycol.

FMN binds to subunit NDUFV1 at the top of the peripheral arm, where NADH is oxidized and the electrons are fed into the chain of Fe-S clusters that runs vertically through the peripheral arm, with one “off-pathway” Fe_2_S_2_ cluster (N1a). The Fe-S cluster chain ends at cluster N2 which transfers electrons to the Q substrate in the Q-binding cavity near the junction between the peripheral and membrane arms. Arg154 of the NDUFS2 subunit, which is close to N2, is modified to a dimethyl arginine, as also observed in the cryoEM structures of ovine *(13)*, mouse *(22)* and *Y. lipolytica* complex I *(12)*. Most likely the dimethyl arginine modulates the redox potential of the N2 cluster *(33)*.

Subunit NDUFA9 binds one molecule of NADPH. A non-covalently attached NADPH in this position has been found in mammalian, yeast and plant complex I, but its function, if any, is unclear. Most likely it does not participate in electron transfer because of its long distance of ∼37 Å to the nearest Fe-S cluster (N2) *(1)*. Studies in which NADPH binding is blocked indicate that it is required for complex I assembly and stability, rather than for catalytic activity *(34-36)*.

The membrane arm of *Ct*-complex I contains one phosphopantetheine group as a cofactor bound to the second copy of subunit NDUFAB1 (NDUFAB1β) at the tip of the membrane arm. In addition to the cofactors, we identified 17 phospholipids and 14 detergent molecules in the cryoEM map of the DDM-solubilized complex I (**Fig. 2A-B**). In the LMNG-solubilized complex, we identified 32 phospholipids and 3 detergent molecules (**Fig. 2C-D**). No densities for NADH or Q were visible, as both samples were prepared and plunge-frozen without added substrates.

### Two different forms of *C. thermophilum* complex I

To inspect the cryoEM structures for conformational variability and heterogeneity, we performed 3D Variability Analysis (3DVA) in cryoSPARC *(37)* (see Methods). 3DVA of the DDM-solubilized complex indicated a flexing and slight twisting motion of the peripheral arm relative to the membrane arm (**Fig. S2, movie S1**), suggesting a flexible hinge between the two arms. 3DVA of the LMNG-solubilized complex likewise indicated flexibility in the same region. However, the twisting motion of the peripheral arm was much more pronounced than in DDM and accompanied by a major reorganization of the map density in the hinge region (**Fig. S3, movie S2**). 3DVA separated the *Ct*-complex I particles into two different populations, suggesting distinct conformations. Repeated 3DVA of the LMNG dataset with a mask around the hinge region improved the separation of the two particle populations and finally resolved two clusters of 153,568 (87.5 %) and 21,989 (12.5 %) particles (**Fig. S3, movie S3**). Both clusters were 3D-refined separately, yielding maps of 2.39 Å and 2.75 Å resolution for the major and minor cluster, respectively. Model building then produced structures of *Ct*-complex I in two different conformations. We refer to these two conformations as form 1 (major cluster) and form 2 (minor cluster) rather than as open and closed, because no significant opening or closing of the angle between two arms was evident. The transition between the two forms of *Ct*-complex I is characterized by a twisting rather than a tilting motion of the peripheral arm relative to the membrane arm. The transition concurs with substantial changes of the complex I structure in the hinge region, the Q-binding cavity, and the E-channel.

Loops of the Q-binding cavity subunits ND1, NDUFS2 and NDUFS7 (referred to as ND1, PSST and 49-kDa in the bovine complex) rearrange in the transition from form 1 to form 2 (**Fig. 3A**). The loop linking TMH 5 and 6 of subunit ND1 between the Q_s_ and Q_d_ site includes a number of highly conserved residues, in particular Glu211 and Glu223 (see **table S2**). In form 2, Glu211^ND1^ and Glu223^ND1^ form salt bridges with Arg128 and Arg132 of subunit NDUFS7. In form 1, the TMH5-6 loop of ND1 moves out of the Q-binding cavity, and both salt bridges break. The rupture of these two salt bridges is also observed in ovine complex I *(13)* (**Fig. S4A**) and is likely to be a key element in the complex I mechanism. Site-directed mutagenesis indicated a significant decrease in Q reductase activity when residues in the loop region containing Arg128 and 132 of NDUFS7 were targeted *(38)*.

**Fig. 3.**
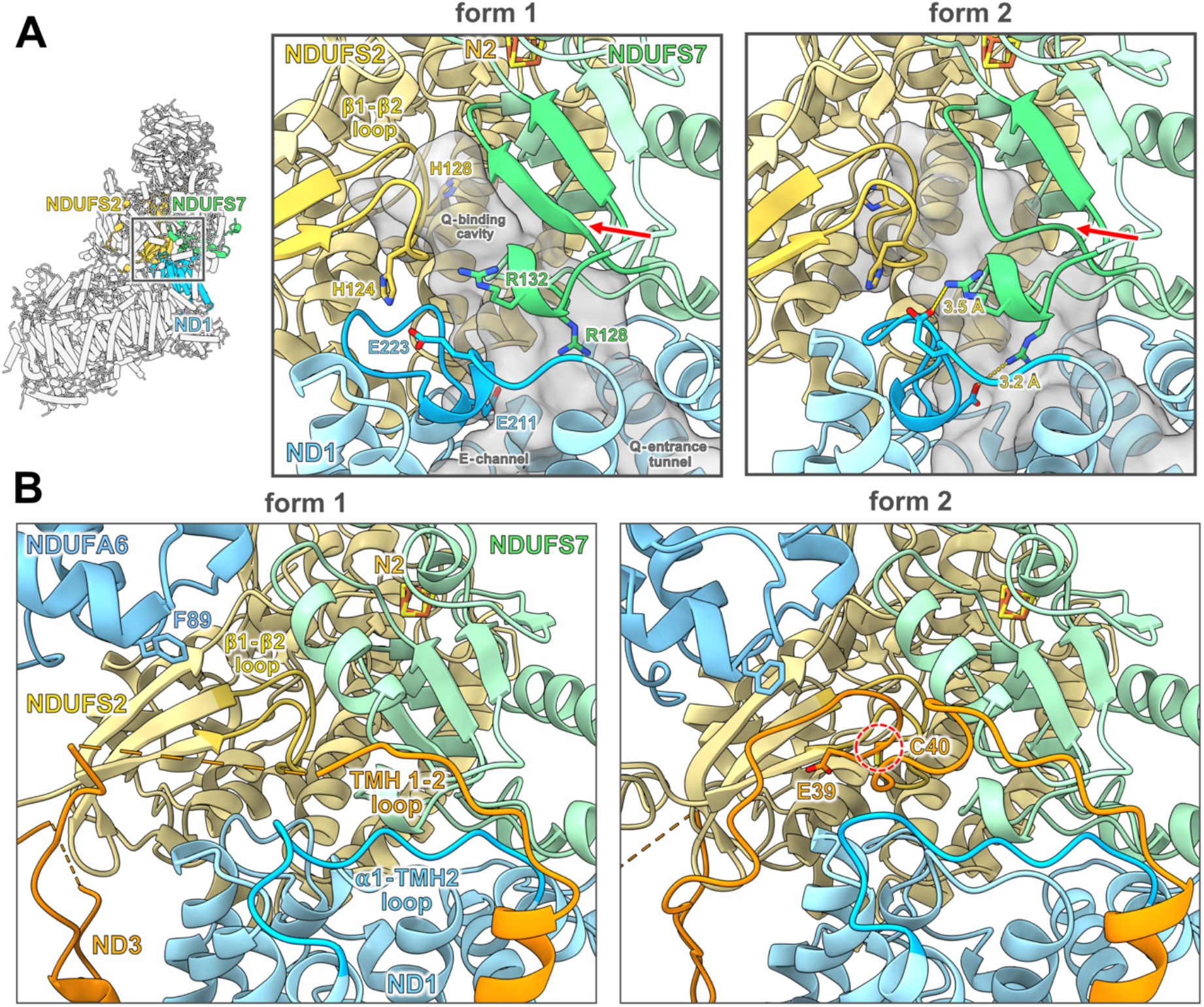
Loops near the Q-binding site reorganize in the 1-to-2 transition. **(A)** In form 2, Glu211^ND1^ and Glu223^ND1^ establish salt bridges with Arg132^NDUFS7^ and Arg128^NDUFS7^. Loop residues 99 to 102 of subunit NDUFS7, which in form 1 are in a β-strand conformation (red arrow), protrude into the Q-binding cavity. The β1-β2 loop of subunit NDUFS2 with its many conserved residues (such as His124 and 128) reorganizes. Helix 3 of subunit NDUFS7 is omitted for clarity. **(B)** The TMH1-2 loop of subunit ND3 with conserved residues Glu39 and Cys40 is disordered in form 1, but becomes ordered and is resolved near the Q_d_ site in form 2, probably supported by the ND1 loop connecting transverse helix α1 to TMH2 (indicated as α1-TMH2 loop) and accessory subunit NDUFA6. Mutation of NDUFA6 residues near the ND3 TMH1-2 loop in *Y. lipolytica* (e.g. Phe89) cause a substantial decrease in complex I activity *(45)*.

The loop between the first and second β-strand of subunit NDUFS2 (known as the β1-β2 loop) is part of the Q_d_ site and includes two conserved histidines (His124 and 128) that are critical for ubiquinone reductase activity of complex I *(39, 40)*. The β1-β2 loop reorganizes in the 1-to-2 transition. In form 1, His128 extends into the Q-binding cavity, but retracts from it in form 2 (**Figs. 3A** and **S4B**). In form 2, loop residues 99 to 102 of subunit NDUFS7 are in an extended conformation and protrude into the Q-binding cavity, but in form 1 they arrange into a β-strand and retract from the Q-binding cavity (**Figs. 3A** and **S4A**).

The loop connecting TMH1 and 2 of subunit ND3 in the membrane arm interacts with peripheral arm subunits that contribute to the Q-binding cavity. In form 1, the TMH1-2 loop is disordered, but in form 2 it is well-ordered and appears to be locked near the NDUFS2 core subunit and accessory subunit NDUFA6 (**Fig. 3B**). In this ordered state the conserved Cys40 in the loop comes close to the Q-binding cavity (**Fig. S5**). The exact role of this loop is under debate, but it clearly is important. Modification of the conserved cysteine has a major impact on complex I activity *(21, 41, 42)*. In the human complex, mutations in this loop result in Leigh syndrome and progressive mitochondrial disease *(43, 44)*. In the 1-to-2 transition, the adjacent ND1 loop connecting transverse helix α1 and TMH2 rearranges and appears to keep the TMH1-2 loop of subunit ND3 in the ordered and defined position seen in form 2 (**Figs. 3B** and **S4C-D**). In addition, site-directed mutagenesis of residues in accessory subunit NDUFA6 close to the TMH1-2 loop of ND3 causes a substantial decrease in complex I activity *(45)* (**Fig. 3B**). We propose that locking the TMH1-2 loop of ND3 in this position by the adjacent subunits ND1 and NDUFA6 is an essential part of the mechanism in mitochondria. Locking of the TMH1-2 loop by subunit NDUFA6 appears to be triggered by the twisting motion of the peripheral arm during the 1-to-2 transition.

We used NADH:Q_1_ oxidoreductase measurements to ascertain that our *Ct*-complex I preparations were active. At 30 °C, the LMNG-solubilized complex showed an activity of ∼10 µmol min^-1^ mg^-1^, which increased to ∼33 µmol min^-1^ mg^-1^, if the measurements were performed at 50 °C (**Fig. S6A**), in line with the thermostability and thermophilic adaptation of *Ct-*complex I. Notably, the activity traces showed no measurable lag phase (**Fig. S6C-D**), which is characteristic of the deactive state as it reverts to the active state. The activity traces closely resemble those for the active ovine complex I *(13)*. Thus, we conclude that *Ct*-complex I in our preparations is predominantly in the active state.

### Internal water molecules rearrange during the conformational transition

Density modification of the cryoEM maps *(46)* reduced noise and improved the overall resolution to the point where water molecules could be built into the density with confidence. We modelled 2,649 water molecules in form 1 of *Ct*-complex I, 1,593 in form 2, and 1,219 in the DDM-solubilized complex.

In the center of the membrane arm, hydrophilic residues coordinate numerous internal water molecules creating an internal aqueous passage (**Fig. S7**). The hydrophilic residues are positioned at discontinuous helix regions of the membrane core subunits. The Q-binding cavity and the aqueous passage through core subunits ND2, ND4 and ND5 are joined via the E-channel, which also contains numerous coordinated water molecules. In form 1, a non-hydrated region in the E-channel at TMH3 of subunit ND6 breaks the aqueous passage, separating the Q-binding cavity from the chain of water molecules in the membrane arm (**Fig. 4**). The peptide geometry of TMH3^ND6^ deviates from that of a regular α-helix and forms a π-bulge at the conserved residue Tyr77 roughly in the middle of the helix. In form 2, the C-terminal half of TMH3^ND6^ on the matrix side that includes the conserved residue Phe85, rotates around its axis by almost 180°, and the π-bulge reverts to a regular α-helix (**Fig. 4A**). Sidechain movements near the TMH3^ND6^ π-bulge (here referred to as the π-gate) create a small internal cavity between TMH3^ND6^, TMH2^ND3^ and TMH3^ND4L^ that becomes hydrated, creating a continuous aqueous passage within the E-channel (**Fig. 4B**). A set of conserved sidechains at the π-gate reorganizes in the 1-to-2 transition, resulting in a rearrangement of water molecules (**Fig. 4B, movie S4**). Leu64 in TMH2^ND3^ rotates by about 180°, enlarging the internal cavity. In form 1, the conserved residues Glu152 and Glu201 of subunit ND1 both form hydrogen bonds to one and the same water molecule. Upon the transition to form 2, TMH4^ND1^ bends near Tyr151^ND1^. The tyrosine sidechain rotates by ∼180° into the E-channel between Glu152^ND1^ and Glu201^ND1^, breaking the hydrogen bond between the two glutamates. Glu152^ND1^ swings towards the π-gate and forms a new hydrogen bond with Asp67 of TMH2^ND3^. On the distal side of the π-gate, rearrangements of the conserved glutamates Glu30^ND4L^, Glu66^ND4L^ and Glu153^ND2^ upon the 1-to-2 transition additionally contribute to a continuous water chain through the π-gate.

**Fig. 4.**
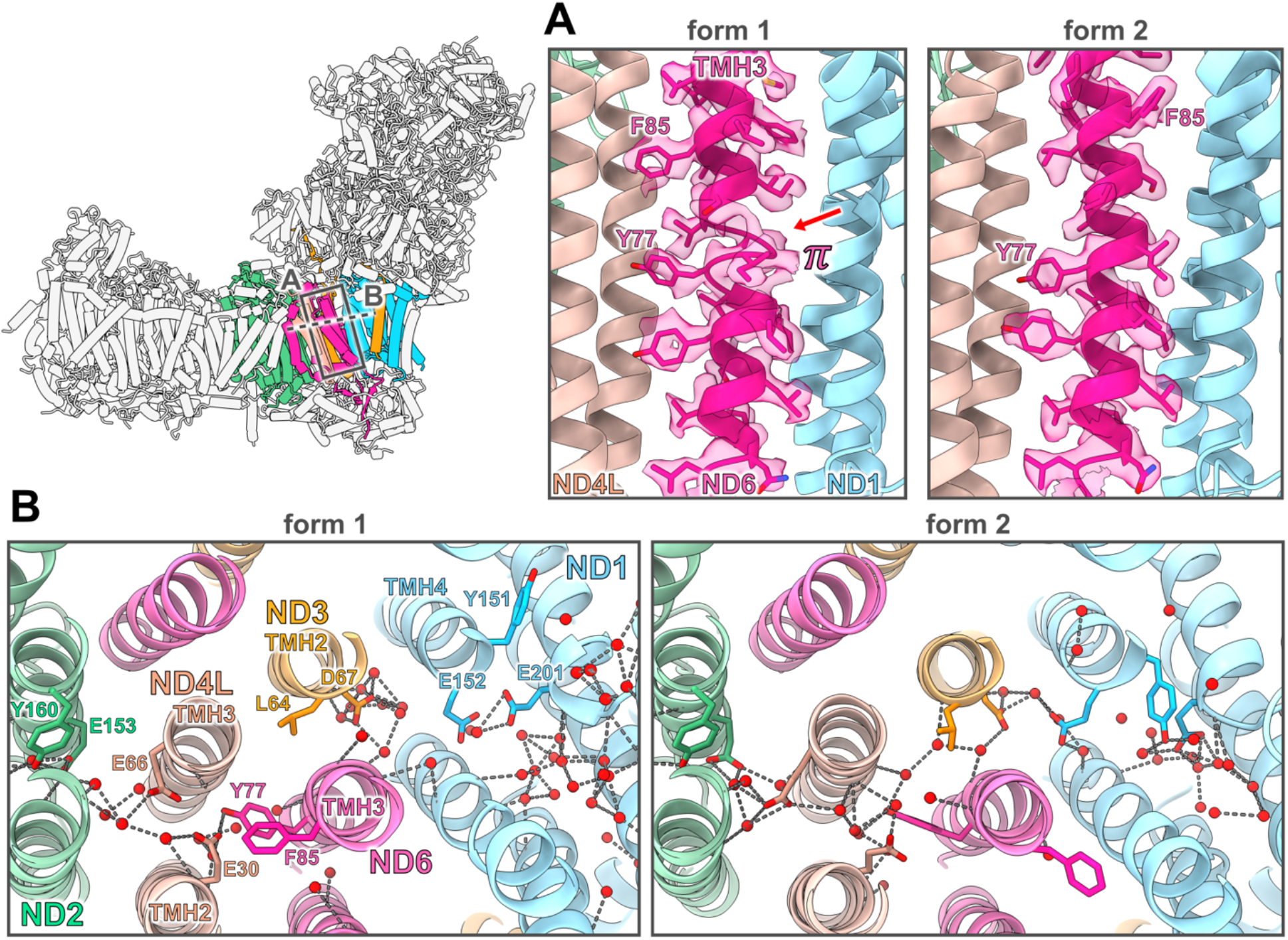
Conformational changes and water reorganization at the π-gate in the 1-to-2 transition. **(A)** In form 1 of *Ct*-complex I, TMH3 of subunit ND6 forms a π-bulge near Tyr77 (red arrow). Upon the transition to form 2, the helix twists by almost 180° at Phe85 and the helix stretches to form a regular α-helix. **(B)** Apart from the reorganization of the π-bulge in TMH3^ND6^ (here referred to as the π-gate), several conserved residues in the E-channel reorganize, creating an interconnected chain of water molecules between the E-channel and the hydrophilic passage within the membrane arm (see **movie S4**).

In form 1 of *Ct-*complex I, the C-terminus of peripheral arm subunit NDUFA9 extends along the hinge connecting the two arms. It folds into two short α-helices that sit like a latch on top of the loop connecting TMH4^ND6^ and TMH3^ND6^ that forms the π-gate (**Fig. 5**). We refer to this C-terminal structure of subunit NUDFA9 as the NDUFA9 latch. In this form 1, Ser360 in the NDUFA9 latch interacts with the conserved Glu98^ND6^. In form 2, the latch retracts from the TMH3-4 loop of ND6 and becomes mostly disordered. This rearrangement coincides with a stretching of TMH3^ND6^ and the reorganization of the π-bulge back into an α-helical conformation (**Fig. 5A**), which is crucial for the water chain through the π-gate.

**Fig. 5.**
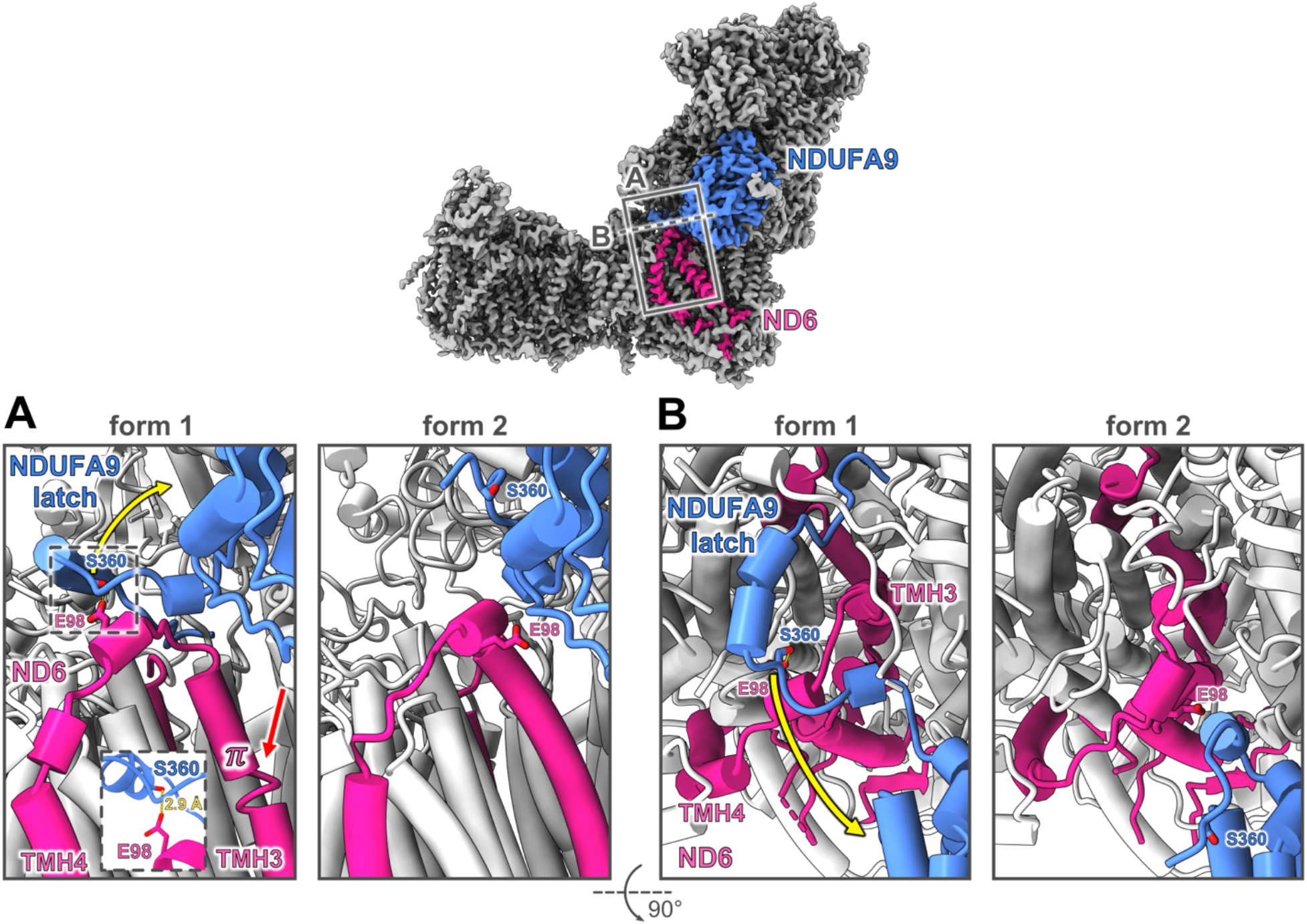
Reorganization of the NDUFA9 latch. **(A)** Side and **(B)** top view of subunit ND6 and NDUFA9. In form 1 of *Ct*-complex I, the C-terminus of subunit NDUFA9 extends along the hinge region of the peripheral and membrane arm and forms a latch across the TMH3-4 loop of subunit ND6. Glu98^ND6^ is coordinated by Ser360^NDUFA9^ (inset). In this conformation TMH3^ND6^ is slightly compressed and forms a π-bulge (red arrow). In form 2, the NDUFA9 latch retracts from the TMH3-4 loop and becomes mostly disordered while TMH3^ND6^ stretches and the π-bulge disappears. Yellow arrows indicate the relocation of Ser360^NDUFA9^ in the 1-to-2 transition.

Focused 3D-refinement improved the resolution of the membrane arm and enabled us to model a total of 136 water molecules along the entire hydrophilic passage (**Fig. S7**). Of the three antiporter-like core subunits ND5, ND4 and ND2, only ND5 at the tip of the membrane arm seems to form an aqueous connection to the IMS, as also observed in *Y. lipolytica (12)* and the ovine complex I *(13)*. The transition between the two forms of complex I did not appear to be associated with a rearrangement of water molecules or conformational changes beyond the π-gate in the membrane arm.

Most of the water molecules in *Ct*-complex I were modelled in the peripheral arm. Apart from the hinge region and the Q-binding cavity, these water molecules did not rearrange in the 1-to-2 transition. As in the ovine complex *(13)*, none of the ∼1,700 water molecules in the hydrophilic peripheral arm were found directly between adjacent Fe-S clusters.

### A bridge between the peripheral and membrane arm

Accessory subunit NDUFA5 is part of the peripheral arm. In *C. thermophilum* this subunit is larger than in any other known complex I structure. It interacts with subunit ND2 in the membrane arm, forming a protein bridge (here referred to as NDUFA5 bridge) between the peripheral and membrane arms that has not been observed before (**Fig. S8**). Remarkably, C and N-terminal sequence extensions of core subunits ND3 and NDUFS2 tether these subunits to the NDUFA5 bridge (**Fig. 6**). The N-terminal extension of NDUFS2 (here referred to as tether 1) is bound to the surface of the NDUFA5 bridge. It directly precedes the β1-β2 loop, which is well-known to be involved in Q binding and reduction *(39, 40)*. Tether 1 therefore establishes a direct connection between the β1-β2 loop and the NDUFA5 bridge. In the transition between the two forms, tether 1 reorganizes (**Fig. 6**). In form 1, tether 1 is exposed on the surface of the membrane arm, extends across the tip of the NDUFA9 latch and interacts with the latch via a salt bridge between Arg114^NDUFS2^ and Glu368^NDUFA9^ close to the β1-β2 loop (**Fig. 6**, inset a). In form 2, tether 1 relocates, enabling a rearrangement of Arg114^NDUFS2^. Also, the NDUFA9 latch retracts from its position at the hinge between the two arms and the salt bridge between Glu368^NDUFA9^ and Arg114^NDUFS2^ breaks (**Fig. 6**).

**Fig. 6.**
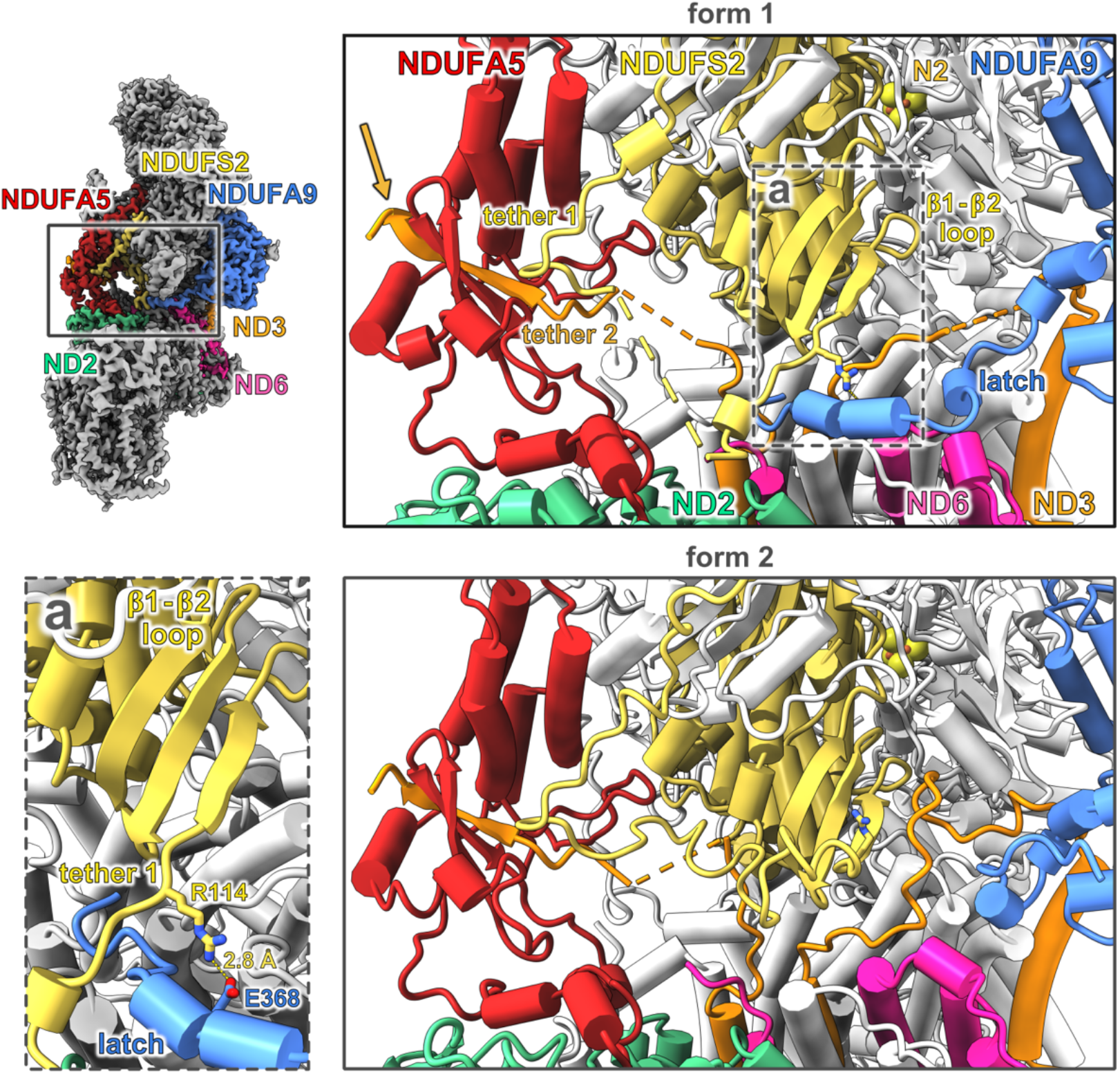
Restructuring of the *Ct*-complex I arm interface. Subunit NDUFA5 interacts with core subunit ND2, forming a protein bridge. Extensions of subunit NDUFS2 and ND3 bind to the NDUFA5 bridge via tether 1 and 2. The C-terminus of tether 2 forms a β-strand in the NDUFA5 bridge (orange arrow). Tether 1 at the N-terminus of subunit NDUFS2 is directly linked to the β1-β2 loop. It is attached to the surface of the NDUFA5 bridge and rearranges upon the 1-to-2 transition. In form 1, tether 1 lies above the tip of the C-terminus of subunit NDUFA9, which is part of the NDUFA9 latch structure. A salt bridge is established between Glu368^NDUFA9^ and Arg114^NDUFS2^ close to the β1-β2 loop (inset a). In form 2, tether 1 rearranges, the salt bridge breaks and the NDUFA9 latch retracts from subunit ND6 and the arm interface. Subunit NDUFAB1α and NDUFA6 are omitted for clarity.

The C-terminal extension of subunit ND3 is enveloped by subunit NDUFA5 and remarkably contributes a β-strand to the β-sheet of the NDUFA5 bridge (**Fig. 6**, orange arrow). This β-strand and TMH3 of subunit ND3 are directly linked by another peptide tether (tether 2), which rearranges and attaches to subunit NDUFS2 in the 1-to-2 transition (**Fig. 6**).

### Detergent effects in *C. thermophilum* complex I

In the DDM-solubilized complex, the ND1 subunit has a 40-residue extension joining TMH 6 and 7 that is absent in other complex I structures. In the DDM structure, the ND1 extension is ordered and forms a hook (the ND1 hook) extending to the detergent belt (**Fig. 7**, see **fig. S9** for a sequence alignment), which is notably thinner (∼25 Å) in this region than around the remainder of the membrane arm. Lipids in this region of the matrix leaflet are clearly tilted, with their acyl chains oriented almost parallel to the membrane plane (**Fig. 2B**). Several amphipathic helices from core and accessory subunits, including NDUFA9, NDUFA12, NDUFS8 or NDUFS7 are located near the matrix leaflet at subunit ND1 and are likely involved in orienting the lipids in this region *(16)*. The distortion of the lipid bilayer reduces the local membrane thickness, which may facilitate access to the Q-entrance tunnel, framed by TMH1, 6, and the transverse helix α1 of subunit ND1 *(16)*. The ND1 hook in *Ct*-complex I may maintain a thinner lipid bilayer near the Q-entrance tunnel, perhaps to improve accessibility for the Q substrate. Presumably this loop is unstable in the LMNG-solubilized complex, due to some difference in shape, size or surface properties of the detergent micelle.

**Fig. 7.**
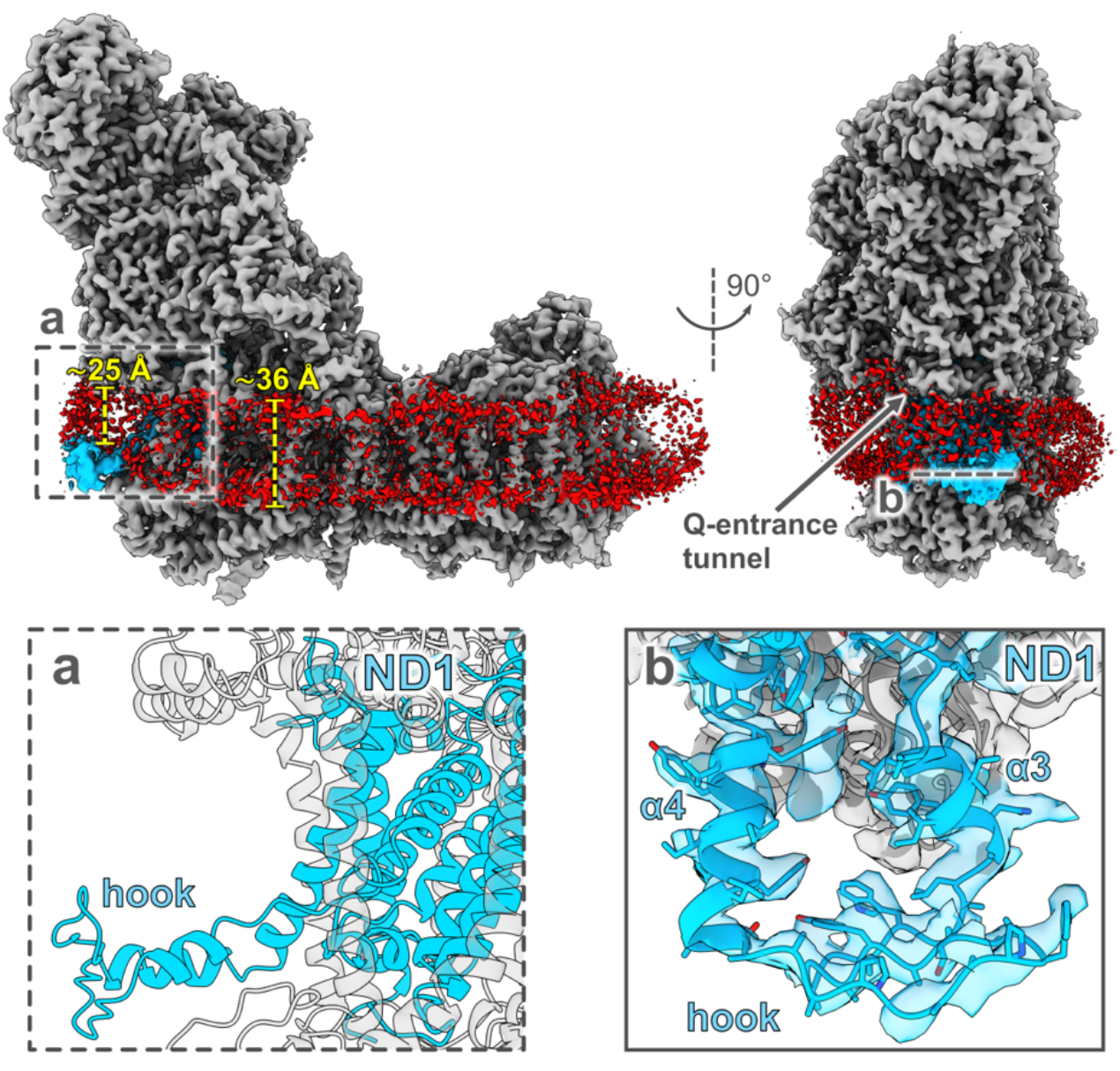
The ND1 hook. The core subunit ND1 (blue) of *Ct*-complex I forms a hook (inset a) that extends onto the DDM detergent belt (red) near the Q-entrance tunnel. In the vicinity of the ND1 hook the thickness of the detergent micelle is reduced from 36 Å to 25 Å. Inset b: CryoEM map density of the ND1 hook with fitted side chains.

Surprisingly, we did not observe the two forms 1 and 2 of *Ct*-complex I in the DDM-solubilized complex. In this detergent, all particles assumed a single conformation, which closely resembled form 1 in LMNG, except that the ND1 loop connecting TMH5 and 6 forms a short α-helix and is shifted towards the N2 cluster (**Fig. 8**). We identified two molecules of DDM near the Q-binding cavity (**Figs. 8A** and **S10**). One of these (DDM-1) binds at the Q-entrance tunnel, as also observed in the DDM-solubilized *Yl*-complex I *(47)*. The hydrophilic headgroup of DDM-1 sits near the Q_s_ site, coordinated by Asp121 of subunit NDUFS7, and its hydrophobic acyl tail extends into the Q-entrance tunnel. DDM-2 is bound in the E-channel with its headgroup coordinated by the conserved residues Glu211 and Glu236 of subunit ND1, and pointing into the Q-binding cavity. The movement of the ND1 loop towards N2 appears to trigger a reorganization of the β1-β2 loop of NDUFS2. The β1-β2 loop protrudes into the Q_d_ site, where it would interfere with the coordination of the Q head group by the conserved Tyr177 in subunit NDUFS2 *(48)* (**Fig. 8**, inset b).

**Fig. 8.**
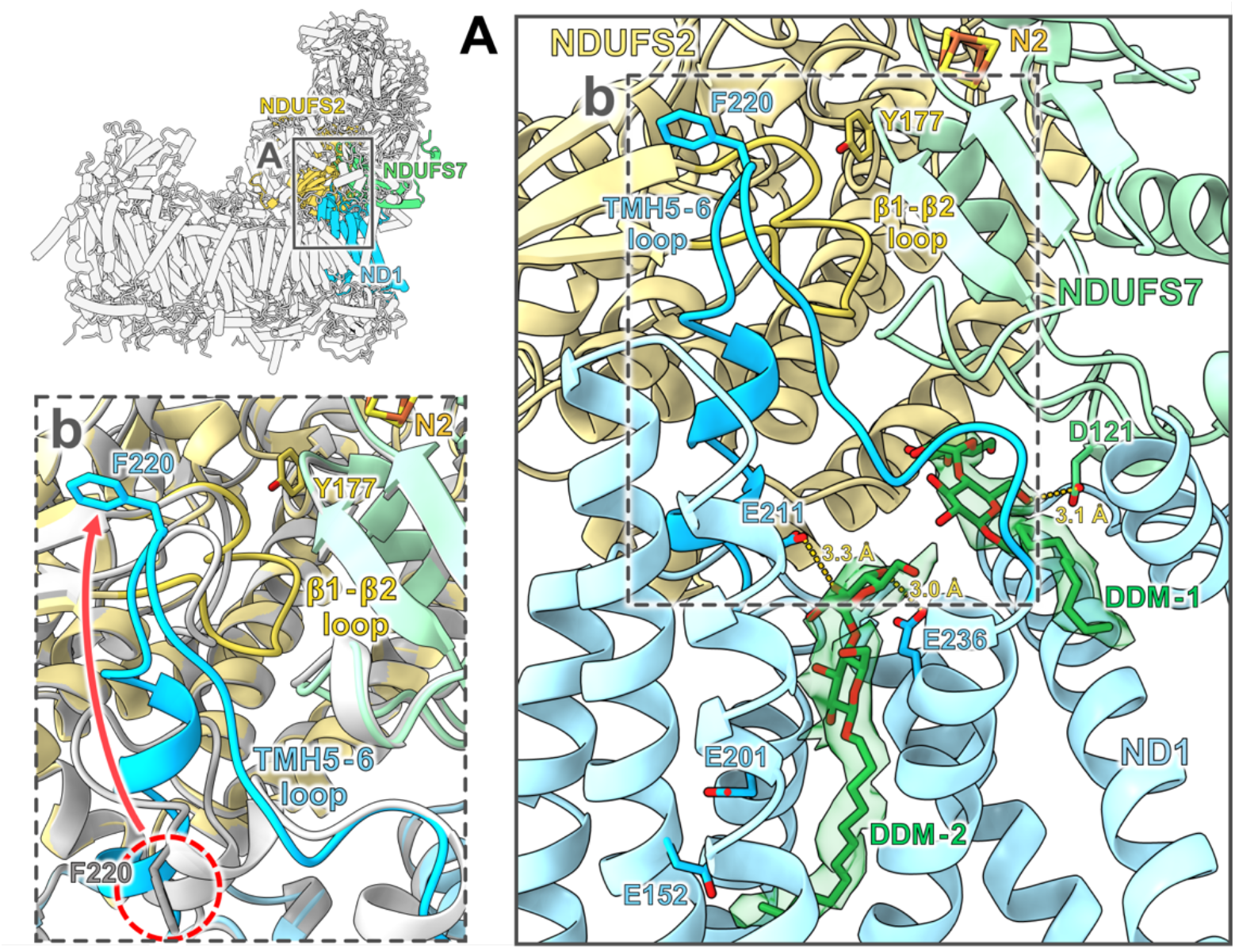
DDM bound in *Ct*-complex I. **(A)** In DDM-solubilized *Ct*-complex I, one DDM molecule is bound in the Q-entrance tunnel (DDM-1) with its head group near the Q_s_ site, coordinated by Asp121^NDUFS7^. A second DDM molecule binds in the E-channel (DDM-2) with its head group coordinated by Glu211^ND1^ and Glu236^ND1^. DDM molecules are shown with EM density. **Fig. S10** shows their position in the Q-entrance tunnel and E-channel. Inset b: Superposition of *Ct*-complex I in DDM (colored by subunit) and in LMNG in form 1 (grey). Overall, the two structures are very similar except that in the DDM structure the TMH5-6 loop of subunit ND1 is shifted towards N2 and forms a partial α-helix. The TMH5-6 loop reorganization is accompanied by a rearrangement of the β1-β2 loop of subunit NDUFS2, which protrudes into the Q_d_ site in DDM-solubilized *Ct*-complex I. Red arrow: Movement of Phe220^ND1^ by ∼16 Å.

Consistently, our activity measurements of *Ct*-complex I solubilized in DDM show only weak Q_1_ reduction activity, compared to complex I in LMNG (**Fig. S6A+B**, *p* < 0.01 between complex I in LMNG and DDM at 30 and 50 °C, without rotenone). Rotenone inhibits the oxidoreductase activity of *Ct*-complex I significantly. However, at 30 °C the activity of DDM-solubilized *Ct*-complex I is so low that rotenone does not inhibit the reaction further (**Fig. S6B**, DDM buffer, 30 °C). In agreement with the observed occupation of the Q-binding cavity this indicates a clear inhibitory effect of DDM on the activity of *Ct*-complex I. Presumably, at 50 °C the inhibitory effect of DDM compared to rotenone is diminished by the decreased binding affinity of the detergent in the Q-binding cavity at elevated temperatures (**Fig. S6B**, DDM buffer, 50 °C). Notably, the oxidoreductase activity increases significantly when DDM-solubilized *Ct*-complex I is transferred to a buffer containing LMNG (**Fig. S6B**, LMNG buffer). The quick recovery indicates that inhibition by DDM is reversible.

## Discussion

CryoEM of mitochondrial complex I from the thermophilic eukaryote *C. thermophilum* revealed that *Ct*-complex I undergoes a transition between two different conformations, which we refer to as form 1 and form 2. The transition is characterized by a twisting motion of the peripheral arm relative to the membrane arm and by a substantial restructuring of the hinge region, the Q-binding cavity, and the E-channel (**Fig. 9**). We are aware that two different conformations have been described also of the mammalian *(13, 22, 23)* and plant complex I *(24)*. In mammals, the two conformations are referred to either as open and closed *(13)* or as the deactive and active state of complex I *(22)*.

**Fig. 9.**
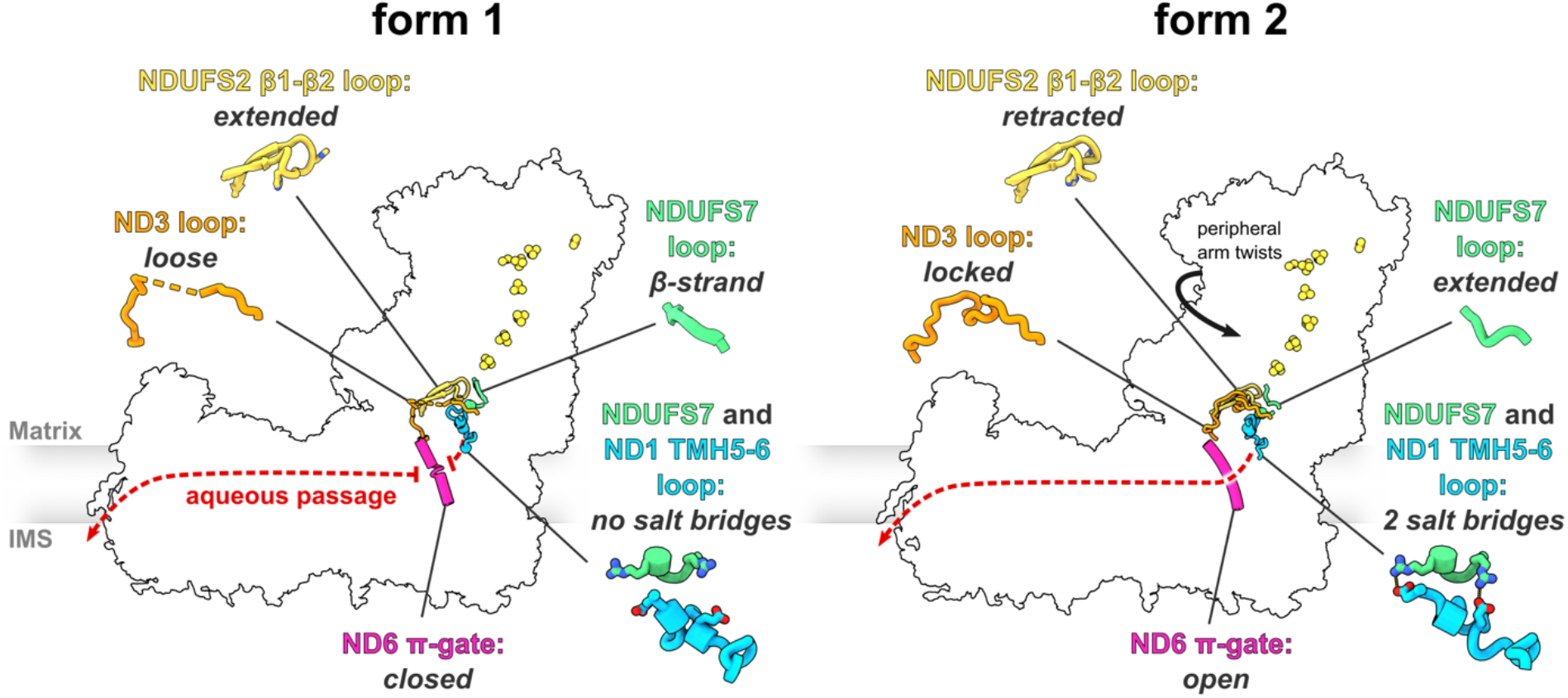
Conformational changes in *C. thermophilum* complex I. The conformational transition between forms 1 and 2 entails a twisting motion of the peripheral arm relative to the membrane arm, as well as rearrangements in the Q-binding cavity and the E-channel. The Q-binding cavity is located close to the interface between the two arms at the end of the chain of iron-sulphur clusters (orange and yellow spheres). Loops of subunits ND1, ND3, NDUFS2 and NDUFS7 in and around the Q-binding cavity undergo substantial changes. The π-gate connects the E-channel and Q-binding cavity to the aqueous passage inside the membrane arm (dashed red arrow). In form 1, the π-gate formed by TMH3^ND6^ is closed, breaking the aqueous passage. In form 2, the π-gate opens as TMH3^ND6^ regularizes into an α-helix, resulting in a continuous aqueous passage from the Q-binding cavity along the membrane arm.

NADH:Q_1_ activity measurements with the *Ct*-complex I did not indicate a lag phase that would be expected of the deactive state. As the sample used for cryoEM grid preparation was treated identically, we conclude that our structures show the active state of *Ct*-complex I. Kampjut and Sazanov drove the ovine complex I into the deactive state by incubating it at 37 °C in the absence of substrates. The structure of the complex deactivated in this way closely resembled the open conformation (our form 1), but indicated a ∼ 40° tilt of TMH4 of subunit ND6. The tilt of this helix is thought to arrest complex I in the deactive state, which presumably represents an off-pathway state of the “open” complex *(13)*. No tilt of TMH4^ND6^ was apparent in any of our *Ct*-complex I structures, consistent with our conclusion that the two forms of *Ct*-complex I are both in an active state.

The open/closed terminology derives from the slightly more open (around 112°) or more closed angle (around 105°) between the two arms in the mammalian complex *(13, 23)*. Since this angle is not noticeably different in the two forms of the *C. thermophilum* complex, the terms open or closed do not apply. The cryoEM structure of plant complex I from *Arabidopsis thaliana* mitochondria *(24)* likewise revealed two conformations, one with a narrower (106°), the other with a slightly wider angle (112°) between the two arms which, in analogy to the mammalian complex, were also referred to as “open” and “closed”. However, neither of them showed the rearrangements in the E-channel or the Q-binding cavity observed in *Ct*-complex I (this work) and in the ovine complex *(13)*. The angle between the membrane and peripheral arm also varies in the bacterial complex I from *Escherichia coli*, for which three different conformations have been be described *(49)*. None of these three conformations indicated a comparable rearrangement in the Q-binding cavity or E-channel. Evidently, both the mitochondrial and bacterial complex I can adopt “open” and “closed” conformations. However, these conformations do not necessarily relate to rearrangements in the Q-binding cavity or the E-channel that are likely to be functionally relevant. The open and closed conformations may simply reflect the intrinsic flexibility of complex I.

Restructuring of the E-channel in the 1-to-2 transition results in the formation of a continuous water chain through the π-gate formed by TMH3^ND6^ of subunit ND6. Similar water and residue reorganizations at the π-gate were also observed in ovine complex I *(13)*. However, the π-gate is closed in the “open” conformation (our form 1), and open in the “closed” conformation (our form 2) (**Fig. S11**), which is potentially confusing. We propose that the two conformations should be referred to simply as form 1 and form 2, until a consensus on their functional relevance has been reached.

The opening of the π-gate in the 1-to-2 transition creates a continuous aqueous passage from the Q-binding site all the way to the antiporter-like subunit ND5, where protons can be expelled into the IMS. The higher resolution of our *Ct*-complex I structure revealed additional water molecules at the open π-gate that contribute to an uninterrupted water chain in form 2 (**Fig. S11**) and suggests that protons can flow easily from the Q-binding sites to the interior of the membrane arm. An essential role of TMH3^ND6^, which contains the π-gate, is consistent with its high degree of sequence conservation *(50)* and with an impaired complex I activity as a result of mutations in this region *(51, 52)*. The formation of a continuous aqueous passage from the Q-binding site to subunit ND5 through the π-gate appears to be a key element of the complex I mechanism.

Substantial rearrangements during the transition between the two forms of complex I also affect the Q-binding cavity. Reorganization of the ND1 TMH5-6 loop, the NDUFS2 (49-kDa) β1-β2 loop and the NDUFS7 (PSST) loop within the Q-binding cavity are implicated in regulating the diffusion of Q intermediates between the binding sites and the ejection or uptake of QH_2_ and Q *(13, 38)* and appear to be well-conserved between mammalian *(13)* and *Ct*-complex I (**Fig. S4**).

The structures of these loops are strikingly similar in the closed ovine complex and in form 2 of the *C. thermophilum* complex I, down to the level of sidechain positions and orientations (**Fig. S4**). By contrast, in form 1 there are slight but clear differences in position and orientation between residues in these core subunit loops of the ovine and *Ct*-complex I. In some cases, they are even disordered and therefore unresolved. In this context it is interesting that decylubiquinone (DQ) binding at the Q_d_ site near Fe-S cluster N2 was observed only in the closed ovine complex I *(13)*, i.e. the equivalent of form 2 of *Ct*-complex I. Presumably, the better-defined conformation of these loops in form 2 is important for locking the Q substrate near N2 for reduction. The ND3 TMH1-2 loop and its essential cysteine adopt a defined and conserved position near the Q_d_ site in the transition to form 2 (**Fig. S4D** and **S5**). Whether this is critical for Q binding, reduction and protonation remains to be investigated.

The structural rearrangements of the Q-binding cavity and E-channel in the open-to-closed transition of ovine complex I are very similar to the rearrangements we observe in the 1-to-2 transition of the *Ct*-complex I and thus appear to be well-conserved (**Fig. S4** and **S11**). Conformations of complex I from the bacterium *Thermus thermophilus* show movements of the peripheral arm that are also linked to rearrangements in the Q-binding cavity and in TMH3 of subunit Nqo10, which is homologous to subunit ND6 of the eukaryotic complex. These rearrangements appeared to be similar although much less pronounced than in the *Ct* or ovine complex. The authors nevertheless assumed that they might correspond to the open and closed conformation of the ovine complex I *(53)*. *T. thermophilus* complex I does not undergo an active-to-deactive transition *(54)*, therefore it can be excluded that any of its conformations represent the deactive state. The fact that complex I from bacteria, mammals, and now also fungi undergoes a transition that is linked to very similar rearrangements in the Q-binding cavity and in the E-channel, in particular at TMH3^ND6^, leads us to the assumption that such a transition represents a principal and conserved feature of the complex I mechanism.

*Ct*-complex I solubilized in the detergent DDM did not undergo the 1-to-2 transition or any of the structural rearrangements observed in LMNG. NADH:Q_1_ activity measurements with *Ct*-complex I indicated strong inhibition by DDM. A similar effect has been reported for the detergent Triton X-100 *(55)*. We discovered two DDM molecules bound in the Q-entrance tunnel and the E-channel. In line with our activity measurements, we conclude that DDM bound to this site prevents substrate binding and inhibits enzymatic activity. As the 1-to-2 transition was not apparent in this detergent, we propose that DDM inhibits the transition. Ovine complex I retains its ability to undergo the transition with the inhibitor rotenone bound in the Q-binding cavity *(13)*. However, rotenone does not have a flexible hydrophobic tail, whereas the acyl chain of one DDM molecule extends into the E-channel and obstructs it. Since the E-channel is a site of major reorganization in the 1-to-2 transition, its obstruction by a bound inhibitor likely contributes to preventing this transition.

In terms of its overall structure, *Ct*-complex I resembles that from other eukaryotes closely, but it also has some interesting aspects that so far appear to be unique. The enlarged subunit NDUFA5 forms a protein bridge between the peripheral and membrane arms that has not been seen in any other complex I structure. The NDUFA5 bridge seems to stabilize the connection between the two arms and might therefore help to maintain structural integrity at elevated temperatures. Recent structures of plant complex I revealed a protein bridge between the two arms formed by the subunits NDUFAB1α, NDUFA6 and the plant-specific ferredoxin-like subunit C1-FDX. This bridge is on the opposite side of the membrane arm, as seen from the peripheral arm, and may have a regulatory function *(24)*. The NDUFA5 bridge in *Ct*-complex I is connected to the core subunits NDUFS2 and ND3 by two polypeptide tethers. Tether 1 is formed by subunit NDUFS2 and connects the NDUFA5 bridge directly with the β1-β2 loop of NDUFS2, which rearranges in the 1-to-2 transition and is a key element in the Q reductase activity *(39, 40)*. We assume that tether 1 imposes a dragging force on the β1-β2 loop and might be an element for regulating its function. Tether 2 of subunit ND3 forms a β-strand within NDUFA5, connecting the NDUFA5 bridge directly to TMH3 of subunit ND3. Tether 2 is likely to stabilize the NDUFA5 bridge by forming a β-strand that is incorporated into a β-sheet in the NDUFA5 bridge. Thus, we speculate that the NDUFA5 bridge combines stabilizing and regulatory functions in *Ct*-complex I.

The structural features described above have not been found in complex I from other species, including that from *Y. lipolytica*, the only other structure of a fungal complex I. Sequence alignments suggest that non-thermophilic relatives of *C. thermophilum*, such as *Neurospora crassa* or *Podospora anserina*, have similar extensions in subunit NDUFA5 and ND1 (**Fig. S8C** and **S9**). In the absence of complex I structures from these organisms it is difficult to say whether these particular features represent adaptations of *Ct*-complex I to increased growth temperatures or have developed at some point in fungal evolution.

In *Ct*-complex I, the C-terminus of subunit NDUFA9 extends into the hinge between the peripheral and membrane arms. It forms a latch that rearranges in the 1-to-2 transition and interacts closely with subunits NDUFS2 and ND6. In form 1, the latch establishes a salt bridge with tether 1 of subunit NDUFS2 close to the β1-β2 loop. As this salt bridge breaks in form 2, we assume that the latch participates in the rearrangement of the critical β1-β2 loop. In form 1, the NDUFA9 latch extends across the TMH3-4 loop of subunit ND6. In the transition to form 2, the latch retracts from this position, while the π-bulge in TMH3^ND6^ reverts to a regular α-helix. The π-bulge is likely to form or disappear in response to some compression or stretching force on TMH3^ND6^, which in *Ct*-complex I appears to be exerted by the NDUFA9 latch via the TMH3-4 loop. Structures of mammalian and plant complex I do not show a NDUFA9 latch. In these organisms, its role must be taken over by some other element of the convoluted complex I structure. In *Y. lipolytica* the NDUFA9 latch is present and notably disordered under turnover conditions *(12)*, consistent with a regulatory role in fungal complex I.

In the transition between form 1 and 2 of complex I, the peripheral arm twists, and this movement triggers conserved rearrangements in the Q-binding cavity and the formation of a water chain through the π-gate, enabling proton and charge transfer from the E-channel to the aqueous passage of the membrane arm. The cryoEM structures of *Ct*-complex I in two different conformations provide strong support for a common mechanism of coupling electron transfer in the peripheral arm to proton translocation in the membrane arm.

## Materials and Methods

### Isolation of mitochondria from *Chaetomium thermophilum*

Wildtype *Chaetomium thermophilum* (La Touche) *var. thermophilum* was obtained from DSMZ, Braunschweig, Germany (No. 1495) and grown as described *(25)* with slight modifications. Mycelium was grown on LB-agar plates at 52 °C for 2 days. 500 ml of CCM medium were inoculated with freshly scraped off and chopped mycelium in a 1 l Erlenmeyer flask. After 1 day at 52 °C and 100 rpm in a rotary shaker, the submerged cultures were shredded in a blender for 10 s and used to inoculate 1 l CCM in 5 l Erlenmeyer flasks. Cultures were incubated at 52 °C for ∼18 h at 100 rpm. For harvest, cultures were strained through a metal sieve (180 µm pore size) and excessive liquid was squeezed out with a silicone spatula. Cell pellets were stored at -20 °C until use.

Mitochondria were isolated from ∼100 g of mycelium. All subsequent steps were performed at 4 °C or on ice. Cells were resuspended in isolation buffer (50 mM Hepes/NaOH pH 7.8, 350 mM sorbitol, 1 mM EGTA, 1 mM PMSF) and lysed by sonication with a Branson Models 250 device at 25 % amplitude, in 2 cycles of 2 min each, with 3 s pulses and 2 s pause times. The cell lysate was centrifuged at 1500 x g for 5 min and 4000 x g for 5 min to remove cell debris. Mitochondria were pelleted by centrifugation at 12,000 x g for 15 min. To improve purity, mitochondria were resuspended in isolation buffer and the last two centrifugation steps repeated. Mitochondria were finally resuspended in resuspension buffer (10 mM Hepes/NaOH pH 7.8, 350 mM sorbitol, 1 mM EGTA) and the concentration of mitochondrial protein was measured by a Bradford assay. The concentration of the resuspension was adjusted to ∼10 mg/ml with resuspension buffer and mitochondria were snap-frozen in liquid nitrogen and stored at -80 °C until further use.

### Purification of complex I

Purified mitochondria from *C. thermophilum* containing ∼50 mg mitochondrial protein were thawed on ice and centrifuged at 12000 x g for 10 min at 4 °C. Supernatant was discarded and the membrane pellet resuspended either in DDM solubilization buffer (100 mM Hepes/NaOH pH 7.8, 2 mM MgCl_2_, 50 mM NaCl, 3 % [w/v] n-dodecyl β-D-maltoside [DDM]) or in LMNG solubilization buffer (100 mM Hepes/NaOH pH 7.8, 2 mM MgCl_2_, 50 mM NaCl, 2.5 % [w/v] lauryl maltose neopentyl glycol [LMNG]) to a final detergent:protein weight ratio of 2:1. The resuspension was rotated for 30 min at 4 °C. Insoluble material was removed by centrifugation at 21,000 x g for 15 min at 4 °C. The supernatant was filtered (0.22 µm pore size) and loaded onto a POROS GoPure HQ column (Thermo Fisher Scientific) connected to an Äkta Purifier system (GE Healthcare). The column was equilibrated with IEX buffer 50 (30 mM Hepes/NaOH pH 7.8, 2 mM MgCl_2_, 50 mM NaCl) supplemented with 0.015 % [w/v] DDM, if mitochondria were solubilized with DDM, or 0.0015 % [w/v] LMNG, if mitochondria were solubilized with LMNG. After loading the column was washed with IEX buffer 50 supplemented with detergent, until a constant baseline was reached. Complex I was eluted with a linear gradient to 70 % of the high-salt buffer IEX 1000 (30 mM Hepes/NaOH pH 7.8, 2 mM MgCl_2_, 1000 mM NaCl) over 60 min with 1 ml/min flowrate. IEX buffer 1000 was supplemented with detergent in the same way as IEX buffer 50. Fractions containing complex I were concentrated using Amicon Ultra 4 columns with 100,000 molecular weight cutoff and loaded onto a Superose 6 Increase 3.2/300 size-exclusion column (GE Healthcare) connected to an Äkta Ettan system. Complex I was eluted in SEC buffer (20 mM Hepes/NaOH pH 7.4, 100 mM NaCl), supplemented with the same type and amount of detergent as IEX buffer 50. Eluted complex I was used directly for cryoEM specimen preparation.

### Sample vitrification and cryoEM data acquisition

1. µl of purified complex I at a final concentration of 3.6 mg/ml (DDM-solubilized) or 1.5 mg/ml (LMNG-solubilized) was applied onto freshly glow-discharged (15 mA for 45 s in a PELCO easiGlow system) C-flat 1.2/1.3 300 mesh copper grids (Science Services GmbH). Samples were blotted for 4 s with blot force 20 (595 Whatman paper) at 4 °C in 100 % humidity and plunge-frozen in liquid ethane with a FEI Vitrobot Mark IV. CryoEM data were collected automatically using EPU software (Thermo Fisher Scientific) on a Titan Krios G3i microscope at 300 kV, equipped with a K3 detector (Gatan) operating in electron counting mode. Movies were acquired at a nominal magnification of 105,000x, resulting in a pixel size of 0.837 Å. Each movie was recorded for 2.1 s and subdivided into 45 frames. The electron flux rate was set to 15 e^-^ per pixel per second at the detector, resulting in an accumulated exposure of 45 e^-^/Å^2^ at the specimen. An energy filter with a slit width of 30 eV was used and a 70 µm C2 and a 100 µm objective aperture were inserted during acquisition. 1,996 movies at a defocus range of -1 µm to -3.5 µm were collected for complex I solubilized in DDM and 6,570 movies at a defocus range of -0.5 to -3.0 µm for complex I solubilized in LMNG. An overview of the data collection statistics is shown in **table S3**.

### Image processing

Image processing was performed in RELION3 *(56, 57)*, unless otherwise specified. For complex I in DDM, dose-fractionated movies were motion-corrected using the RELION implementation of the MotionCor2 algorithm *(58)* and contrast transfer function (CTF) parameters were estimated with CTFFIND4 *(59)*. 126,638 particles were picked in crYOLO 1.7.3 with a trained model from 20 micrographs *(60)*. Particle coordinates were imported into RELION and were extracted with down-sampling to a pixel size of 3.348 Å. Particles were then subjected to 2D classification, resulting in 24,016 particles, which were selected to reconstruct an Initial model. The Initial model was used as a reference for 3D classification with 102,840 particles, selected from classes of the previous 2D classification. After 3D classification, broken and low-resolution particles were sorted out and the remaining 37,815 particles were re-extracted with a pixel size of 1.0299 Å and a box size of 512 pixels. An additional round of 3D classification was performed. The remaining particles were subjected to 3D refinement followed by two rounds of per-particle CTF refinement with an intermediate Bayesian polishing step. After removal of particle duplicates, the refined particles were transferred to cryoSPARC v3.0.1 *(37)* to perform a non-uniform refinement in combination with per-particle-defocus and per-group CTF parameter refinement *(61)*, which resulted in a map with an overall resolution of 2.77 Å.

For complex I in LMNG, motion correction and CTF estimation were again performed in RELION with implementation of the MotionCor2 algorithm and CTFFIND4, respectively. Micrographs with an estimated resolution below 7 Å were excluded and 1,087,651 particles were picked from the remaining 6,503 micrographs in crYOLO. Particles were extracted with a down-sampled pixel size of 3.348 Å. After 2D classification, 177,728 particles were selected and subjected to an ab-initio reconstruction in cryoSPARC. The ab-initio reconstruction was used as a reference for 3D classification with 996,588 particles, selected from classes of the previous 2D classification. 3D classes with broken. Junk particles were removed and the remaining 199,973 particles were re-extracted with a box size of 540 pixels, giving a pixel size of 0.9765 Å. The re-extracted particles were subjected to 3D refinement, followed by two rounds of per-particle CTF refinement with an intermediate Bayesian polishing step. A 3D classification yielded a class of 175,581 particles that indicated high-resolution features. As 3D refinement yielded a map of 2.55 Å, close to Nyquist frequency, the particles were re-extracted into 588 pixel boxes with a pixel size of 0.837 Å. Particles were again subjected to two rounds of per-particle CTF refinement with an intermediate Bayesian polishing step and 3D refinement, resulting in a map at 2.50 Å resolution. After removal of particle duplicates, the dataset was further processed in cryoSPARC by performing anon-uniform refinement in combination with per-particle-defocus and per-group CTF parameter refinement, which improved the global resolution to 2.39 Å.

*3D variability analysis* (3DVA) was performed in cryoSPARC *(62)* with the non-uniform refined particles and a real-space mask excluding solvent and the detergent micelle. 3DVA was run using three variability components and a low-pass filter resolution of 6 Å. 3DVA with the DDM dataset revealed flexibility at the hinge between the membrane arm and peripheral arm, but no discrete cluster of particles with a different conformation of complex I. 3DVA of the LMNG dataset indicated a heterogeneous distribution of the particles along the latent coordinate for component 2 and resolved a subset of particles with a strongly twisted peripheral arm in relation to the membrane arm. Repeating 3DVA with the LMNG dataset by masking the hinge region between the peripheral and membrane arm of complex I (where the variability was greatest) and decreasing the low-pass filter resolution to 4 Å, resulted in a better separation of particle subsets along component 2. By using the *cluster mode* function in cryoSPARC, the particle subsets were separated into two clusters of 21,989 and 153,568 particles each. Both particle sets were again subjected to non-uniform refinement in cryoSPARC, which resulted in a map of 2.44 Å (153,568 particles) or 2.78 Å resolution (21,989 particles), showing complex I in two distinct conformations (form 1 and form 2). Additional 3D classifications of the refined particles in RELION (without image alignment and increased T values) confirmed the presence of two conformations in the LMNG-solubilized complex I dataset (with similar particle distribution) and delivered only one conformation for the DDM-solubilized complex I.

To further improve the resolution of the complex I membrane arm in the tip region, the non-uniformly refined particles of form 1 and 2 (in LMNG) and the DDM-solubilized complex, were subjected to a local refinement in cryoSPARC by masking the membrane arm. The locally refined particles of the membrane arm of complex I in DDM yielded a map of 2.76 Å resolution. The membrane arm of form 1 and form 2 resulted in maps of 2.47 Å and 2.83 Å resolution, respectively. All resolutions were estimated according to the Fourier shell correlation (FSC) 0.143 cut-off criterion of two independently refined half maps *(63)*. As a final step, one cycle of density modification with *phenix*.*resolve_cryo_em (46)* was carried out with all 6 reconstructed maps, using in each case the unsharpened map, 2 unfiltered half-maps and an estimate of the protein mass as inputs. This yielded a resolution of 2.75 Å at FSC_ref_ = 0.5 for the map of the entire DDM-solubilized complex I and for the membrane arm. The FSC_ref_ gave a resolution of 2.68 Å for form 2 of the entire complex and 2.75 Å for the membrane arm. For form 1, the FSC_ref_ = 0.5 resolution was 2.32 Å for the entire complex and 2.39 Å for the membrane arm. An overview of the processing workflow and the results is given in **figs. S12 - S14**.

### Model building

Initial models for the *C. thermophilum* complex I subunits were built using the SWISS-MODEL server *(64)* on templates 6RFR and 6RFQ *(16)*. Homology models were rigid-body fitted into the cryoEM map of complex I (form 1) using USCF Chimera *(65)*, followed by manual building in Coot *(66)*. The structure model was rigid-body fitted into all other maps and corrected in Coot. Subunit NDUFB2 and NDUFA1 were not annotated in the genome of *C. thermophilum (29, 67)* and were initially modelled as poly-alanine chains. Characteristic residue-specific map features were subsequently used to place matching amino acids. The resulting short sequence fragments were used as query sequences for protein BLAST *(68)* searches in a 6-frame-translation database from the genome of *C. thermophilum*. Missing residues were replaced by the amino acid sequence of the returned hits. Sequences of some subunits did not match map features, which can be attributed to false exon-intron boundaries in the annotated genome. These errors were corrected manually by extending or shortening the exons in the genomic sequences. Models were iteratively refined by using *phenix*.*real_space_refine* in PHENIX *(69, 70)* and manual refinement in Coot. Ligand restraint files were created by PHENIX eLBOW *(71)*. Water molecules were built automatically using segmentation-guided water and ion modelling (SWIM) *(72)*, implemented in the Segger tool of UCSF Chimera. Water molecules were checked and corrected with the “check/delete waters” function in Coot and also placed manually, if densities had nearly spherical shapes, did not clash with other atoms and showed at least 1 hydrophilic interaction with neighboring residues or waters up to a distance of 3.5 Å. Validation checks for model stereochemistry were performed in MolProbity *(73)*. An overview of the modelling statistics is shown in **table S4**. For structural comparison, models were aligned using the Matchmaker tool in UCSF ChimeraX *(74)*. Cavities were calculated and drawn using Hollow *(75)* with a 1.4 Å probe radius. Finalized models were visualized using UCSF ChimeraX *(74)*. Sequence alignments were performed with Clustal Omega *(76)*.

### Protein analysis by MS

Purified complex I samples were reduced with TCEP and cysteines alkylated with IAA. Subsequent proteolytic digests were performed using S-TRAPs (Protifi) according to the manufacturer’s instructions. Peptides were further desalted and purified on C18 SPE cartridges and dried in an Eppendorf concentrator. After solubilization in 0.1 % formic acid (FA) in acetonitrile/water (95/5 [v/v]), samples were subjected to LC-MS/MS analysis on an Ultimate 3000 nanoRSLC (Thermo Fisher) system, equipped with an Acclaim Pepmap C18 trap column (2 cm * 75 µm, particle size: 3 µm; Thermo Fisher) and a C18 analytical column (50 cm * 75 µm, particle size: 1.7 µm; CoAnn Technologies) with an integrated liquid-junction and fused silica emitter coupled to an Orbitrap Fusion Lumos mass spectrometer (Thermo Fisher). Trapping was performed for 6 min with a flow rate of 6 µl/min using loading buffer (98/2 [v/v] water/acetonitrile with 0.05 % Trifluoroacetic acid) and peptides were separated on the analytical column at a flow rate of 250 nl/min with the following gradient: 4 to 48 % B in 45 min, 48 to 90 % B in 1 min and constant 90 % B for 8 min followed by 20 min column re-equilibration at 4% B with buffer A (0.1 % FA in water) and buffer B (0.1 % FA in 80/20 [v/v] acetonitrile/water). Peptides eluting from the column were ionized online using a Nano Flex ESI-source and analyzed in data-dependent mode. Survey scans were acquired over the mass range from 350 - 1400 m/z in the Orbitrap (maximum injection time: 50 ms, AGC (automatic gain control) fixed at 2×10E5 with 120K mass resolution) and sequence information was acquired by a top-speed method with a fixed cycle time of 2 s for the survey and following MS/MS-scans. MS/MS-scans were acquired for the most abundant precursors with a charge state from 2 to 5 and an intensity minimum of 5×10E3. Picked precursors were isolated in the quadrupole with a 1.4 m/z isolation window and fragmented using HCD (NCE (normalized collision energy) = 30 %). For MS/MS spectra, an AGC of 10E4 and a maximum injection time of 54 ms were used and detection was carried out in the Orbitrap using 30K mass resolution. The dynamic exclusion was set to 30 s with a mass tolerance of 10 ppm. Data analysis was performed in Proteome Discoverer (version 2.5) using Sequest HT as database search algorithm for peptide identification. Raw files were recalibrated and searched against the protein database obtained from structure modelling as well as the Uniprot proteome for *Chaetomium thermophilum* (UP000008066; obtained 2020-10-09) and common contaminants. The search space was restricted to tryptic peptides with a length of 7-30 amino acids allowing for up to two missed cleavages and with a minimum of one unique peptide per protein group as well as precursor and fragment mass tolerances of 10 ppm and 0.02 Da respectively. Carbamidomethylation of cysteine was set as a fixed modification and oxidation of methionine was set as variable modification. Inferys rescoring and Percolator nodes were used to estimate the number of false positive identifications and results filtered for a strict target false discovery rate (FDR) < 0.01.

### Activity measurements

Complex I NADH:Q_1_ oxidoreductase activity was measured at 30 °C or 50 °C by monitoring the oxidation of NADH at 340 nm (ε = 6220 M^-1^ cm^-1^) using a Varian Cary 50 UV-Vis spectrophotometer. Measurements were carried out with 1-5 µg/ml purified complex I protein in 20 mM Hepes/NaOH pH 7.4, 50 mM NaCl, 1 mM EDTA, 2 mM NaN_3_ and with either 0.015 % [w/v] DDM or 0.015 % [w/v] LMNG. The reaction was started by addition of complex I protein, after 100 µM 2,3-dimethyoxy-5-methyl-6-(3-methyl-2-butenyl)-1,4-benzoquinone (Q_1_) and 100 µM NADH were already added to the buffer. For complex I inhibition 2 µM Rotenone was added to the reaction mix. Values represent means ± SEM. Statistical analysis was performed using OriginPro 2020b (OriginLab). Statistical significance was determined at the *p* < 0.05 (*), *p* < 0.01 (**) and *p* < 0.001 (***) levels using independent two-tailed two-sample student *t*-test analysis.

## Acknowledgments

We thank Volker Zickermann and Ahmed-Noor A. Agip for critical comments on the manuscript and Özkan Yildiz for valuable assistance with model building. We thank Nikola Kellner and Ed Hurt for their invaluable help in establishing the culture of *C. thermophilum* in our laboratory.

## Funding

This work was funded by the Max Planck Society.

## Author contributions

E.L. purified and characterized complex I, prepared cryoEM grids, acquired and processed cryoEM data, built and analyzed the atomic models and drew the figures. J.M.-C. and J.L. acquired and analyzed MS data. E.L. and W.K. conceived of the study and wrote the manuscript.

## Competing interests

The authors declare that they have no competing interests.

## Data and materials availability

The cryoEM maps have been deposited in the Electron Microscopy Data Bank with accession codes EMD-14797 (complex I, form 1), EMD-14798 (complex I, form 1, only membrane arm), EMD-14794 (complex I, form 2), EMD-14796 (complex I, form 2, only membrane arm), EMD-14791 (complex I in DDM), EMD-14792 (complex I in DDM, only membrane arm). The atomic models have been deposited in the Protein Data Bank under accession codes 7ZMG (complex I, form 1), 7ZMH (complex I, form 1, only membrane arm), 7ZMB (complex I, form 2), 7ZME (complex I, form 2, only membrane arm), 7ZM7 (complex I in DDM), 7ZM8 (complex I in DDM, only membrane arm). The mass spectrometry proteomics data are available via ProteomeXchange with identifier PXD033234. All data needed to evaluate the conclusions in the paper are present in the paper and/or the Supplementary Materials.

## Notes

### Competing Interest Statement

The authors have declared no competing interest.

