## Supplements for "Conformational changes in mitochondrial complex I from the thermophilic eukaryote *Chaetomium thermophilum*"

**This PDF file includes:**

Figs. S1 to S14
Tables S1 to S4
Captions for Movies S1 to S4
References

**Other Supplementary Materials for this manuscript include the following:**

Movies S1 to S4
Table S5

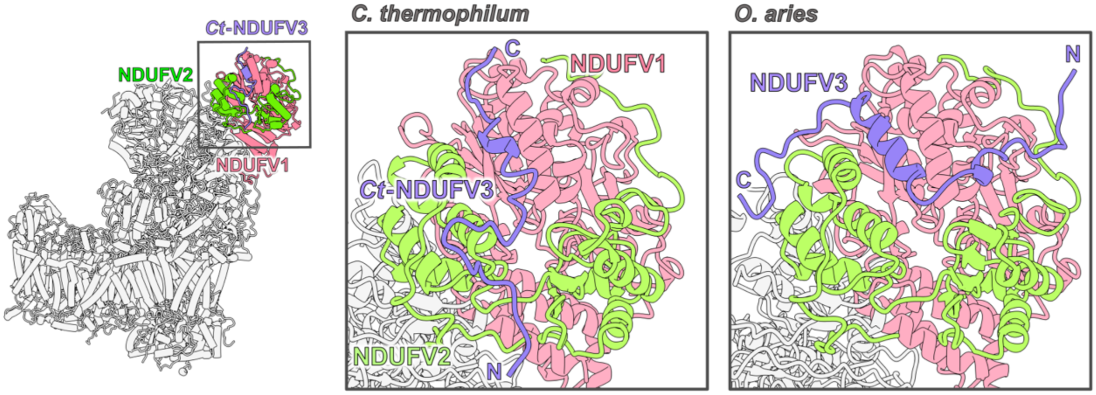


**Fig. S1. Subunit *Ct*-NDUFV3 of *C. thermophilum* complex I.** A 31-residue stretch of subunit *Ct*-NDUFV3 binds in a cleft between subunit NDUFV1 and NDUFV2 on the top of the peripheral arm, where it forms a short α-helix, like its mammalian counterpart (here from *Ovis aries*, PDB: 6ZKR). There was no density for the remaining 349 residues. Since mass spectrometry (MS) detected peptides along the entire length of *Ct*-NDUFV3 (**Table S5**), we conclude that the subunit is mostly disordered. In mammals, subunit NDUFV3 is found in two isoforms (NDUFV3-S [~10 kDa] and NDUFV3-L [~50 kDa]), depending on the tissue, with most of its N-terminal part unstructured (*1, 2*). Not much is known about a potential physiological role of NDUFV3. However, it resembles the 68 kDa fragment of Fat cadherins (*1*), which also binds to complex I at subunit NDUFV2 and promotes oxidative phosphorylation and complex I stability in mitochondria of *Drosophila melanogaster* (*3*). The absence of the 68 kDa fragment results in loss of complex I activity and increased ROS production. It is assumed that NDUFV3-L plays a similar role (*1, 4*). Phylogenetic tree analyses suggest that NDUFV3 is confined to vertebrates, but absent in fungi (*1*). Sequence similarity between NDUFV3 and the newly identified subunit *Ct*-NDUFV3 in our structure is low (7.7 %). However, as the newly identified subunit has a similarly long disordered stretch (although at the C-terminus, rather than the N-terminus) and binds at a nearby site on complex I, we assume that its role is analogous to that of mammalian NDUFV3.

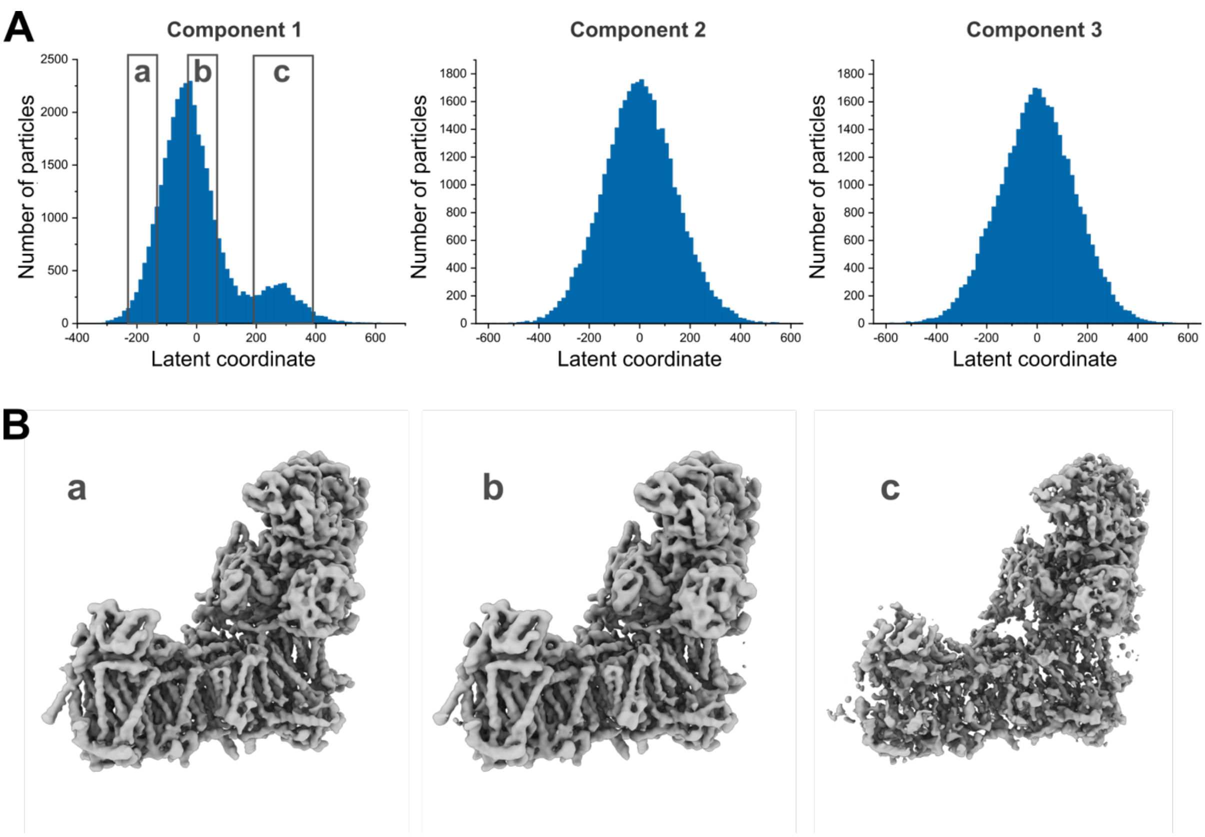


**Fig. S2. 3D variability of complex I in DDM. (A)** One-dimensional histograms of 3DVA for all three variability components. The plots indicate the extent of variability of the particles along each dimension. Component 2 and 3 show a homogeneous distribution and resolved flexibility in the angle between the peripheral and membrane arm (see **movie S1)**. Component 1 resolved a heterogenous distribution and a bimodal split of the particles into two peaks. **(B)** 3D reconstructions of particle clusters along component 1, as exemplary shown (particles reconstructed from the indicated boxes in (A)), revealed that the bimodal split along component 1 is caused by outlier particles, which have lower signal-to-noise ratio or are partially broken (as also described in Punjani and Fleet 2021 (*5*)) and constitute the smaller peak in the histogram (see **movie S1**). Number of particles in the boxes from (A) are 3,630 (a), 12,196 (b) and 4,095 (c). Reconstruction series along each component are shown as movies in **movie S1**.

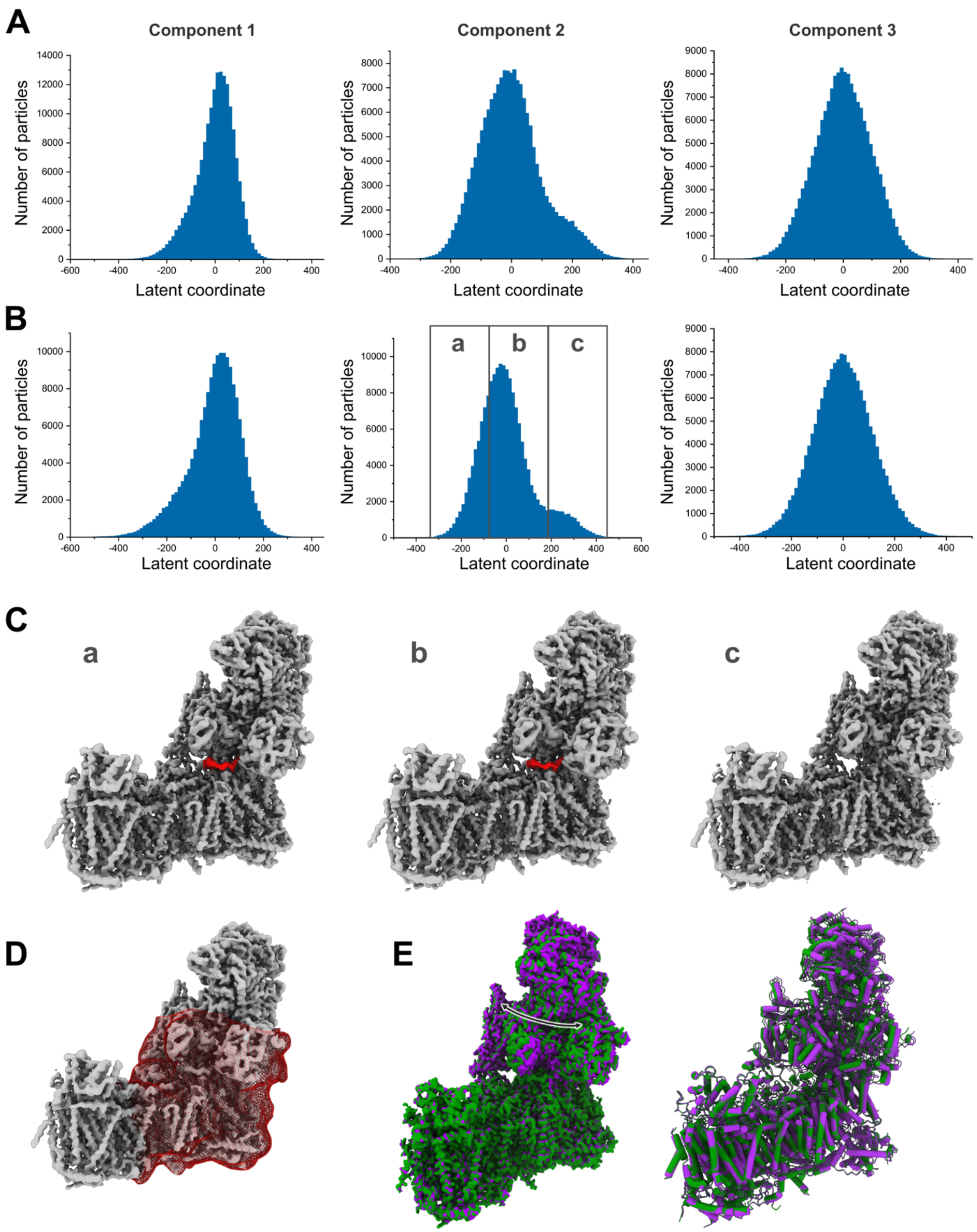


**Fig. S3. 3D variability of complex I in LMNG. (A)** One-dimensional histograms of 3DVA for all three variability components. As for complex I in DDM, component 1 resolved (slight) variability among the particles based on differences in their signal-to-noise ratio or their integrity at the tip of the peripheral arm (see **movie S2**). Component 3 resolved a bending motion between the membrane and peripheral arm of complex I. Component 2 resolved a pronounced twisting motion of the peripheral arm and additionally a heterogenous distribution with two particle populations, representing two different conformations of the complex (see **movie S2**). **(B)** 3DVA was repeated by masking the interface of the peripheral and membrane arm. This resulted in a more distinct separation of both particle populations along component 2. **(C)** Exemplary 3D reconstructions along component 2 of particles from the indicated boxes in (B). Particles of the main peak (box a and b) represent complex I mainly in form 1 and particles of the smaller peak/population (box c) represent complex I mainly in form 2. For the following non-uniform refinement of both particle populations, cluster mode function of cryoSPARC was used and particles were separated into clusters of 153,568 and 21,989 particles. Number of particles in the boxes from (B) are 48,469 (a), 109,050 (b) and 17,624 (c). The C-terminal helix of subunit NDUFA9 (marked in red) shows a clearly visible unfolding in the arm interface upon the transition from form 1 to form 2. Reconstruction series along each component are shown as movies in **movie S2** and **S3**. **(D)** Applied mask for 3DVA as described in (B). **(E)** Overall comparison of the maps (left) and models (right) of complex I in form 1 (green) and form 2 (violet) indicates the twisting motion (~3°) of the peripheral arm.

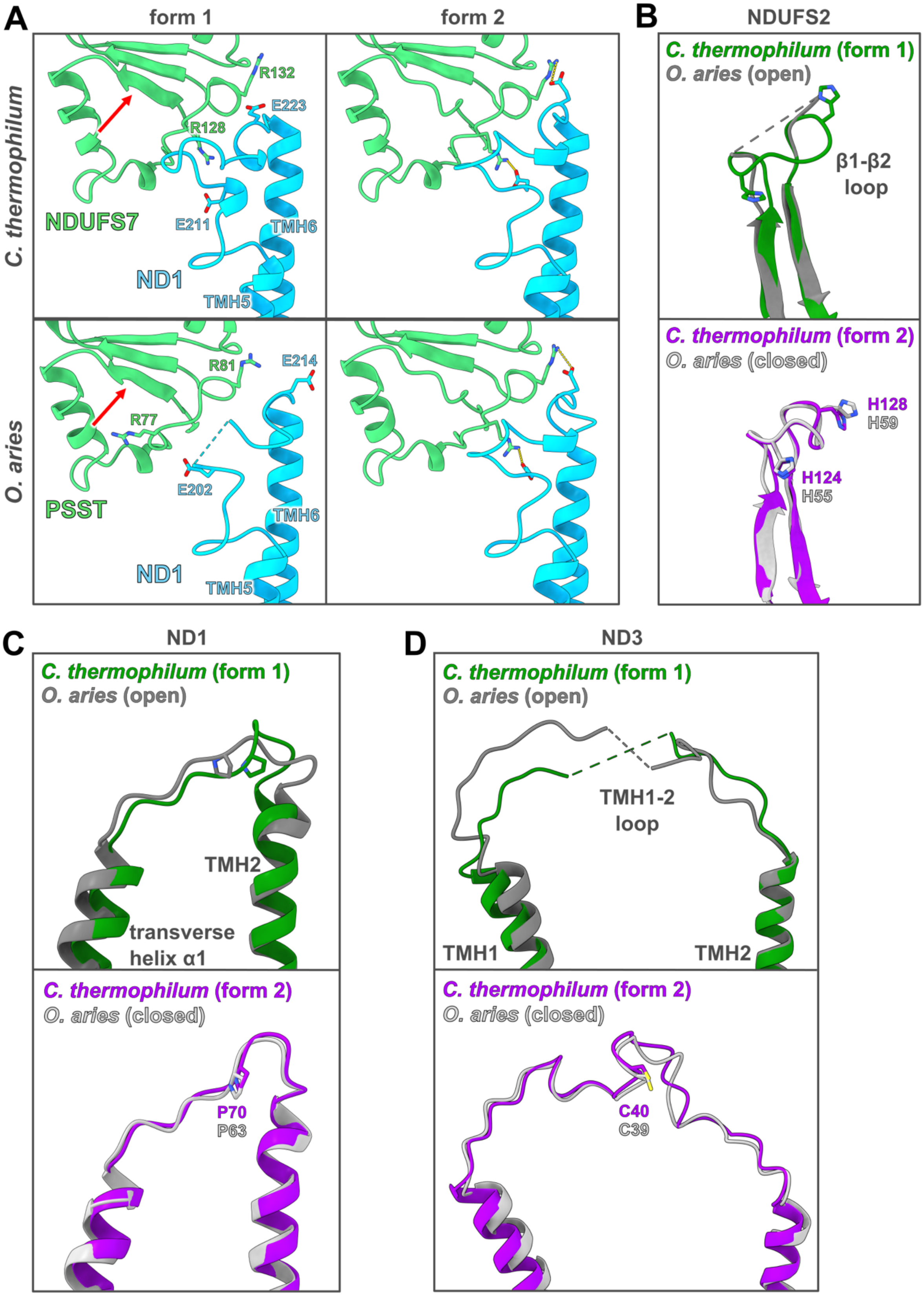

A
form 1
form 2
C. thermophilum
O. aries
R132
E223
R128
NDUFS7
E211
TMH6
ND1
TMH5
R81
E214
R77
E202
PSST
TMH6
ND1
TMH5
B
NDUFS2
C. thermophilum (form 1)
O. aries (open)
β1-β2
loop
C. thermophilum (form 2)
O. aries (closed)
H128
H59
H124
H55
C
ND1
C. thermophilum (form 1)
O. aries (open)
TMH2
transverse
helix α1
C. thermophilum (form 2)
O. aries (closed)
P70
P63
D
ND3
C. thermophilum (form 1)
O. aries (open)
TMH1-2
loop
TMH1
TMH2
C. thermophilum (form 2)
O. aries (closed)
C40
C39

**Fig. S4. Conserved conformational rearrangements in complex I from *C. thermophilum* and *O. aries*.** Upon the transition from form 1 to form 2 (referred to open and closed conformation in ovine complex I, respectively), complex I from *C. thermophilum* and from *O. aries* show common structural rearrangements in several core subunits. Both complexes show especially in form 2 (closed in ovine) high structural similarity. **(A)** The TMH5-6 loop of subunit ND1 rearranges in a conserved way and forms salt bridges to core subunit NDUFS7 in form 2. Two conserved glutamates of the ND1 TMH5-6 loop form salt bridges with conserved arginines of subunit NDUFS7 (referred to as subunit PSST in ovine complex I), respectively. Loop residues 99 to 102 (48 to 51 in ovine complex I) of subunit NDUFS7 extend into the Q-binding cavity in form 2, but form a β-strand in form 1 of complex I (red arrow). **(B)** His128 (His59 in ovine) of the β1-β2 loop of subunit NDUFS2 (referred to as subunit 49-kDa in ovine complex I) extends into the Q-binding cavity in form 1 and is retracted from the cavity in form 2. **(C)** The loop connecting transverse helix α1 and TMH2 of subunit ND1 shows clear rearrangements at Pro70 (Pro63 in ovine) and appears to lock the TMH1-2 loop of ND3 in form 2. **(D)** The TMH1-2 loop of subunit ND3 at the conserved cysteine is unresolved in form 1, but gets resolved in form 2 and the cysteine adopts a defined and conserved position. (PDBs for open and closed ovine complex I: 6ZKP and 6ZKO).

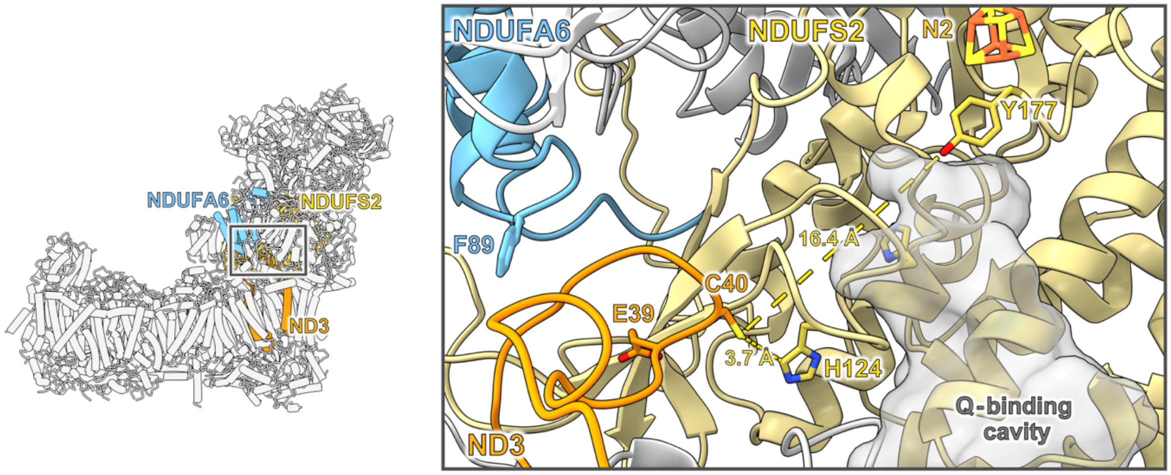


**Fig. S5. Cys40$^{ND3}$ gets close to the $Q_d$ site in form 2 of complex I.** In form 2, the conserved Cys40 in the TMH1-2 loop of subunit ND3 gets close to the Q-binding cavity and is ~3.7 Å and ~16.4 Å away from His124 and Tyr177 of subunit NDUFS2, respectively. His124 and Tyr177 are conserved residues of the $Q_d$ site and crucial for Q binding and reduction (*6-8*). Mutation of NDUFA6 residues near the TMH1-2 loop of ND3 in *Y. lipolytica*, e.g. Phe89, cause substantial decrease in complex I activity *(9)*.

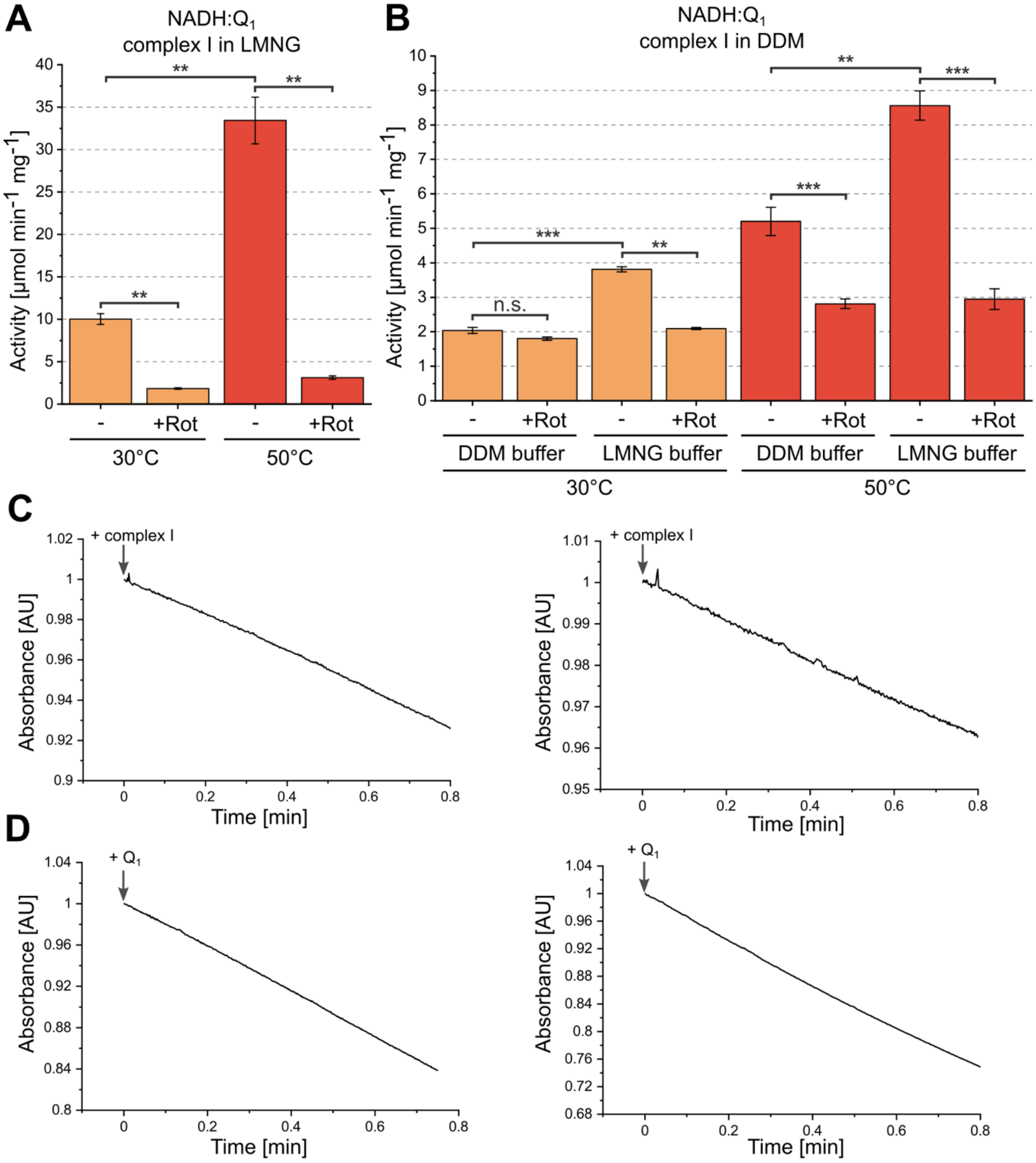


**Fig. S6. Temperature-dependent activity of *Ct*-complex I. (A)** NADH:$Q_1$ oxidoreductase activity of purified complex I solubilized in LMNG without or with inhibitor rotenone (Rot) (n = 3, mean ± SEM). **(B)** NADH:$Q_1$ oxidoreductase activity of purified complex I solubilized in DDM after transfer into buffers with DDM or LMNG as detergent, without or with rotenone (Rot). Notably, activity of complex I in DDM buffer is not significantly different to activity in presence of rotenone at 30 °C. Activity of DDM-solubilized complex I increases significantly when transferred from DDM into LMNG containing buffer (without rotenone) (n = 3, mean ± SEM). Reactions were started by addition of purified complex I to the reaction buffer. **(C)** Representative activity assay traces of complex I solubilized in LMNG at 30 °C (left) and 50 °C (right). Reactions were started by addition of complex I. Activity traces showed no notable lag-phase in reaching the maximum slope. **(D)** For comparison, complex I was pre-incubated with NADH for 1 min before the reaction was started by addition of $Q_1$. Activity traces are similar to (C) (left at 30 °C, right at 50 °C). Note that for measurements at 50 °C less amount of protein was used.

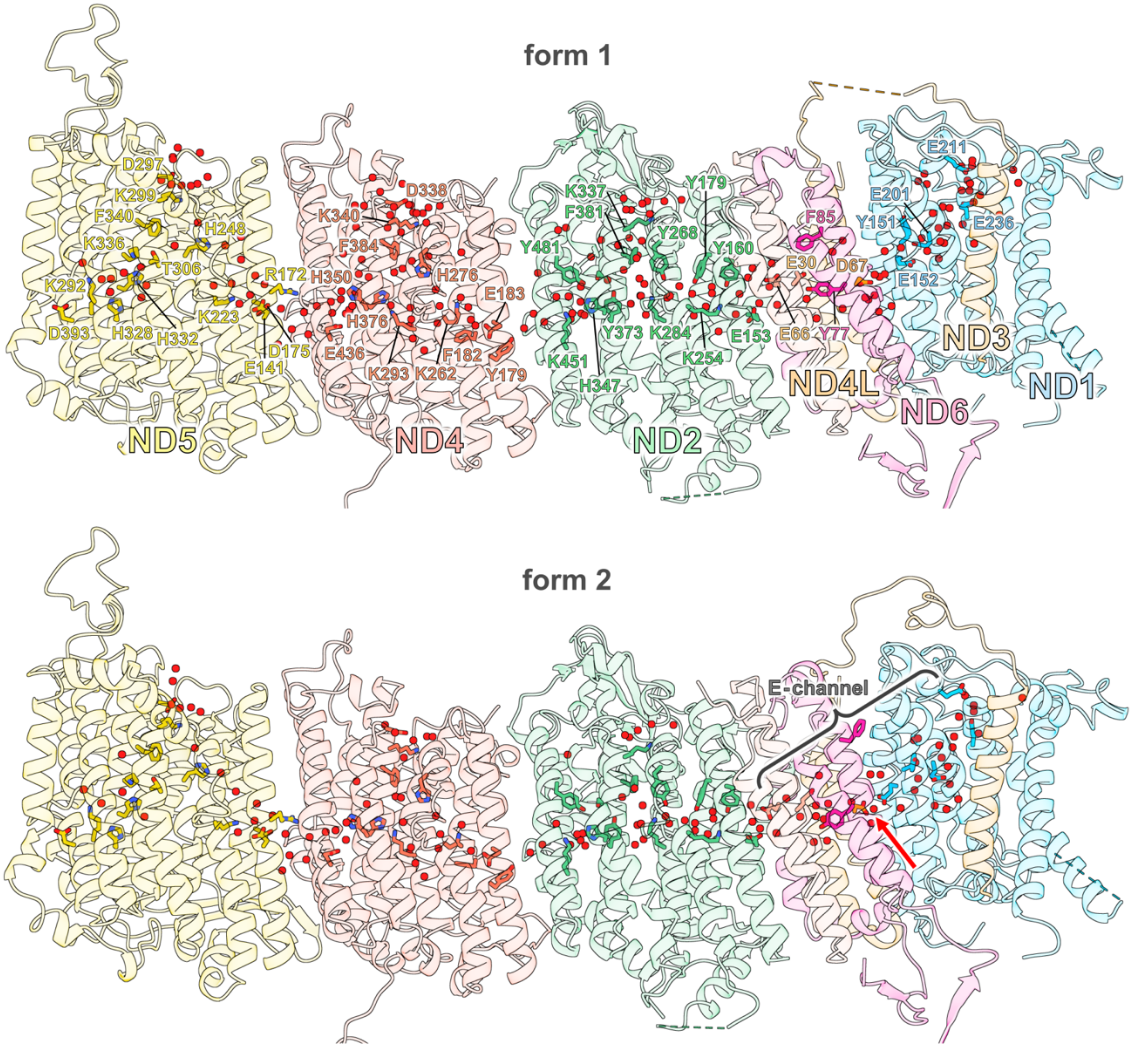


**Fig. S7. Conserved residues and bound water molecules in the membrane arm**. A lane of conserved hydrophilic residues in the membrane arm of complex I is filled with water molecules. Distinct water and residue reorganizations between form 1 and form 2 are only observable along the E-channel (π-gate indicated by red arrow). In form 2, a continuous water wire is formed through the π-gate. Fewer water molecules were modelled in the membrane arm of form 2 due to lower resolution. Only core subunits shown, lateral-helix of subunit ND5 and TMH4 of subunit ND6 omitted for clarity.

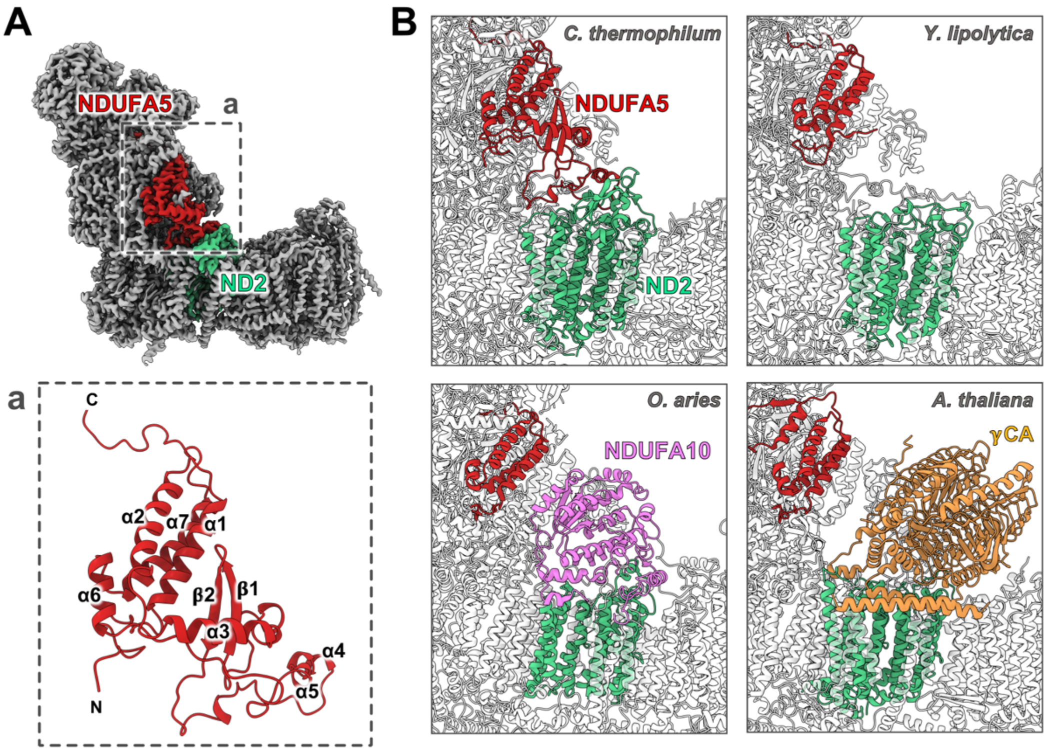


**C**

```
                   α2  α3
C. thermophilum    K-QIPEHSVYRQSVEAVTRHRLALVESVVPEGYDAWVERAKKLLEEHADQFDPTKVGPEG      119
P. anserina        K-AVPEHSVYRQSVEAVTKQRLAHVESVVPPGYKEWAAKAKEILKQEPEKFRTTNTATN-      108
N. crassa          K-QIPEHSLYRQSAEALTKHRLAIVEQYVPDGYDAWQERARKLLEKHKSDLTARQFDGQH      113
Y. lipolytica      QDKHPKDSVYRQSIENLTAHRKQIVEDN--------------------------------      85
H. sapiens         E-EIPKNAAYRKYTEQITNEKLAMVKAE--------------------------------      62
O. aries           G-HIPKNAAYRKYTEQITNEKLSIVKAE--------------------------------      62
A. thaliana        Q-AVPEDEGYRKAVESFTRQRLNVCKEE--------------------------------      71
Consensus              *:.  **:  * .* .:    :

                   β1  β2
C. thermophilum    GEEVQGARAVKVERDGRVFVVSRLPKEEDERVLEWDGEVDEG--------PELEGSRTLE      171
P. anserina        ----EMLGAAKVERDGQVFVVRQLPSEVDMRYQQWDGEVNDG--------PELEGSRTQE      156
N. crassa          A-------RLVEGPDGRAYFIRQMVPPQDWRDVEWDGAVLDPHFSWVQTGEDVVGAVKLE      166
Y. lipolytica      ------------------------------------------------------------      85
H. sapiens         ------------------------------------------------------------      62
O. aries           ------------------------------------------------------------      62
A. thaliana        ------------------------------------------------------------      71
Consensus

                   α4  α5  α6
C. thermophilum    EKIE----------------GGL-EKIFTRRDVKDTVGRVEWEPEPQLTAEQVAELENKI      214
P. anserina        EM-E----------------WHV-KTQFERAE-ALEQKEVEWDPEPQLTTEQIAELENKI      197
N. crassa          DSDKLLELDKIRESDPVAYRQGLRDLGIKMGGVVEDKSPVEWESEPPLSAEQIAEMEARI      226
Y. lipolytica      --------------------------------------------------EVSEVIENKI      95
H. sapiens         --------------------------------------------------PDVKKLEEQL      72
O. aries           --------------------------------------------------PDVKKLEEQL      72
A. thaliana        --------------------------------------------------EDWEMIEKRL      81
Consensus                                                                 :* ::

                   α7
C. thermophilum    GAGLIEEVIQVAEGELKLVDTMVKARVWEPLEEQPRP-GQWEYFERKP------------      261
P. anserina        GAGLIEEVIQVAEGELKLTDTMIESKVWEPLEEPAAE-GQWVAFERTA------------      244
N. crassa          GSGLIEEVVQVAEGELKLVDIMTQARPWEALEEEAPE-GQWTYFERKE------------      273
Y. lipolytica      GAGLIEEVVIQAHEELELAKKMSEWKPWEELEEKPLE-DQWVYFNKKGVE----------      144
H. sapiens         QGGQLEEVILQAEHELNLARKMREWKLWEPLVEEPPA-DQWKWPI---------------      116
O. aries           QGGQIEEVILQAENELSLARKMIQWKPWEPLVEEPPA-SQWKWPI---------------      116
A. thaliana        GCGQVEELIEEARDELTLIGKMIEWDPWGVPDDYECEVIENDAPIPKHVPQHRPGPLPEQ      141
Consensus            * :**::  *. ** *   * :   *    :      :
```

**Fig. S8. Subunit NDUFA5 forms a unique bridge-like structure between the peripheral and membrane arm of complex I. (A)** Subunit NDUFA5 interacts with the matrix side of core subunit ND2. Inset a: Structure of subunit NDUFA5 with secondary structure elements indicated. **(B)** Exemplary comparison of complex I structures from different organisms. Beside NDUFA5, also core subunit ND2 of *C. thermophilum* is slightly larger compared to ND2 structures from other organisms and forms an interaction interface with NDUFA5. No other available complex I structure shows this interaction. A fungal specific adaptation seems unlikely as *Yarrowia lipolytica* shows no similar bridge (PDB: 6RFR). In *Ovis aries* (PDB: 6ZKR), the matrix side of subunit ND2 is occupied by subunit NDUFA10, which has no homologs in plant and fungi *(10)*. In *Arabidopsis thaliana* (PDB: 7ARB) the heterotrimeric γ-carbonic anhydrase (γCA), absent in mammals, fungi and bacteria *(11)*, occupies the matrix side of subunit ND2. **(C)** Sequences of subunit NDUFA5 of complex I from different organisms are compared (sequences only partially shown). Secondary structures of NDUFA5 in *Ct*-complex I indicated above as shown in inset a of (A). Beside *C. thermophilum*, the related fungi *Podospora anserina* and *Neurospora crassa* show similar extensions in the bridge forming parts of subunit NDUFA5. Sequences shown from (Uniprot ID in parentheses): *Chaetomium thermophilum* (G0S8H4), *Podospora anserina* (B2AZ47), *Neurospora crassa* (A0A0B0E3Z9), *Yarrowia lipolytica* (Q6C4W9), *Homo sapiens* (Q16718), *Ovis aries* (W5PNX7), *Arabidopsis thaliana* (Q9FLX7).

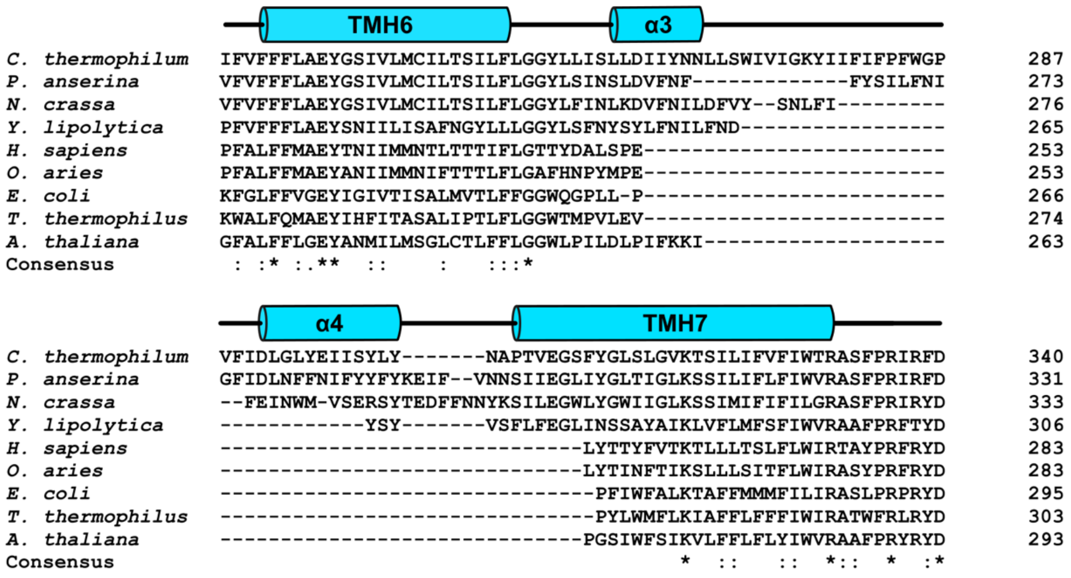


**Fig. S9. Hook-forming extension in subunit ND1.** Sequences of subunit ND1 homologues of complex I from different organisms are compared (sequences only partially shown). Secondary structures of ND1 in *Ct*-complex I indicated above as shown in **fig. 7**, inset b. Beside *C. thermophilum*, the related fungi *Podospora anserina* and *Neurospora crassa* show similar extensions at the hook-like forming part of ND1. Sequences shown from (Uniprot ID in parentheses): *Chaetomium thermophilum* (G1DJA6), *Podospora anserina* (P19041), *Neurospora crassa* (P08774), *Yarrowia lipolytica* (Q9B6E8), *Homo sapiens* (P03886), *Ovis aries* (O78747), *Escherichia coli* (P0AFD4), *Thermus thermophilus* (Q60019), *Arabidopsis thaliana* (P92558).

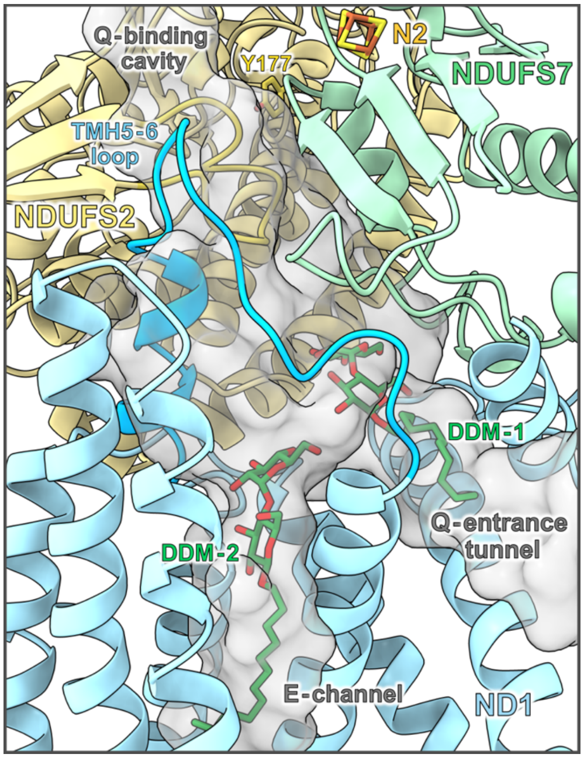


**Fig. S10. Two DDM molecules bind in the Q-entrance tunnel and E-channel of *Ct*-complex I.** In DDM-solubilized *Ct*-complex I, one DDM molecule is bound in the Q-entrance tunnel (DDM-1) and a second one (DDM-2) in the E-channel. Cavity was calculated without the DDM molecules and is shown in grey. See also fig. 8.

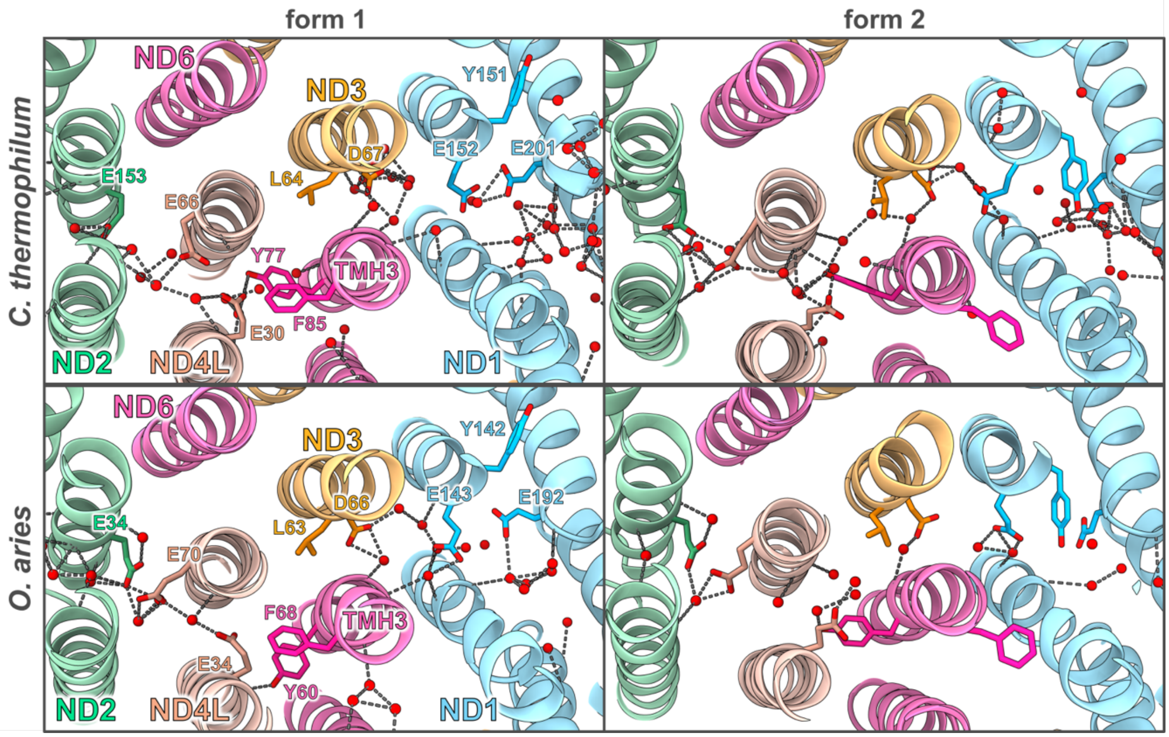


**Fig. S11. The π-gate interrupts the aqueous passage in the membrane arm.** In the E-channel of complex I from *C. thermophilum* (upper row) and *O. aries* (lower row) a π-bulge in $TMH3^{ND6}$ reverts to a regular α-helix in the transition from form 1 to form 2 (referred to as open and closed conformation in ovine complex I). The helix reorganization of $TMH3^{ND6}$ (referred to as π-gate) upon the transition to form 2 is associated with a reorganization of a chain of water molecules passing through the open gate. The structure of form 2 reveals an aqueous passage through the π-gate, in which water molecules and hydrophilic residues are interconnected. H-bonds are shown as dashed lines. (See **movie S4**) (PDBs for open and closed ovine complex I: 6ZKA and 6ZKB).

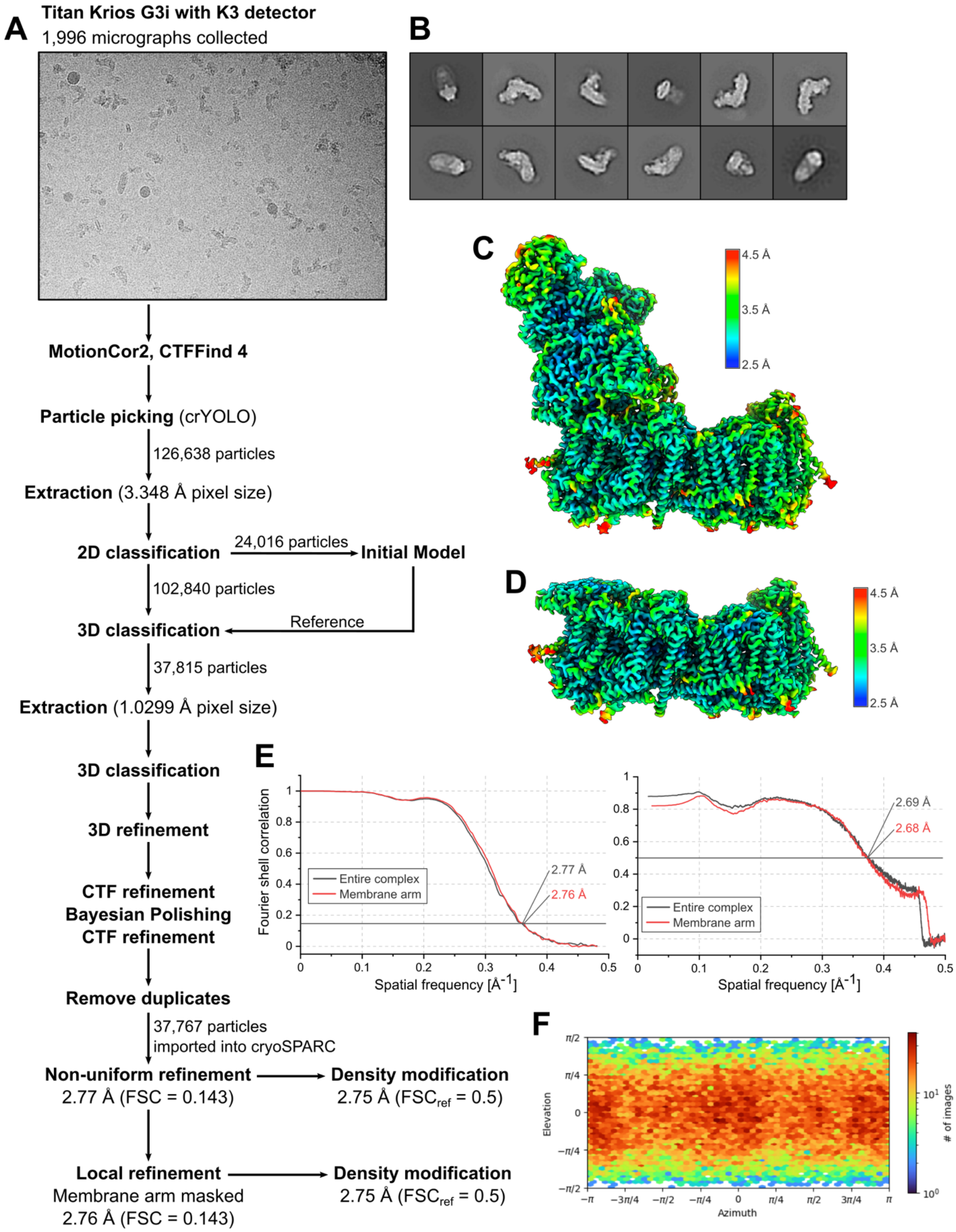


**Fig. S12. CryoEM processing pipeline for complex I in DDM. (A)** Representative micrograph of complex I in DDM and workflow of single-particle cryoEM image processing of complex I solubilized in

DDM. **(B)** Representative 2D class averages. **(C)** Local resolution of the entire complex I map after non-uniform refinement. **(D)** Local resolution of the membrane arm of complex I after local refinement. **(E)** Two-half-map (left) and model-map (right) Fourier shell correlation curves. **(F)** Distribution of orientations over azimuth and elevation angles for particles included in the reconstruction.

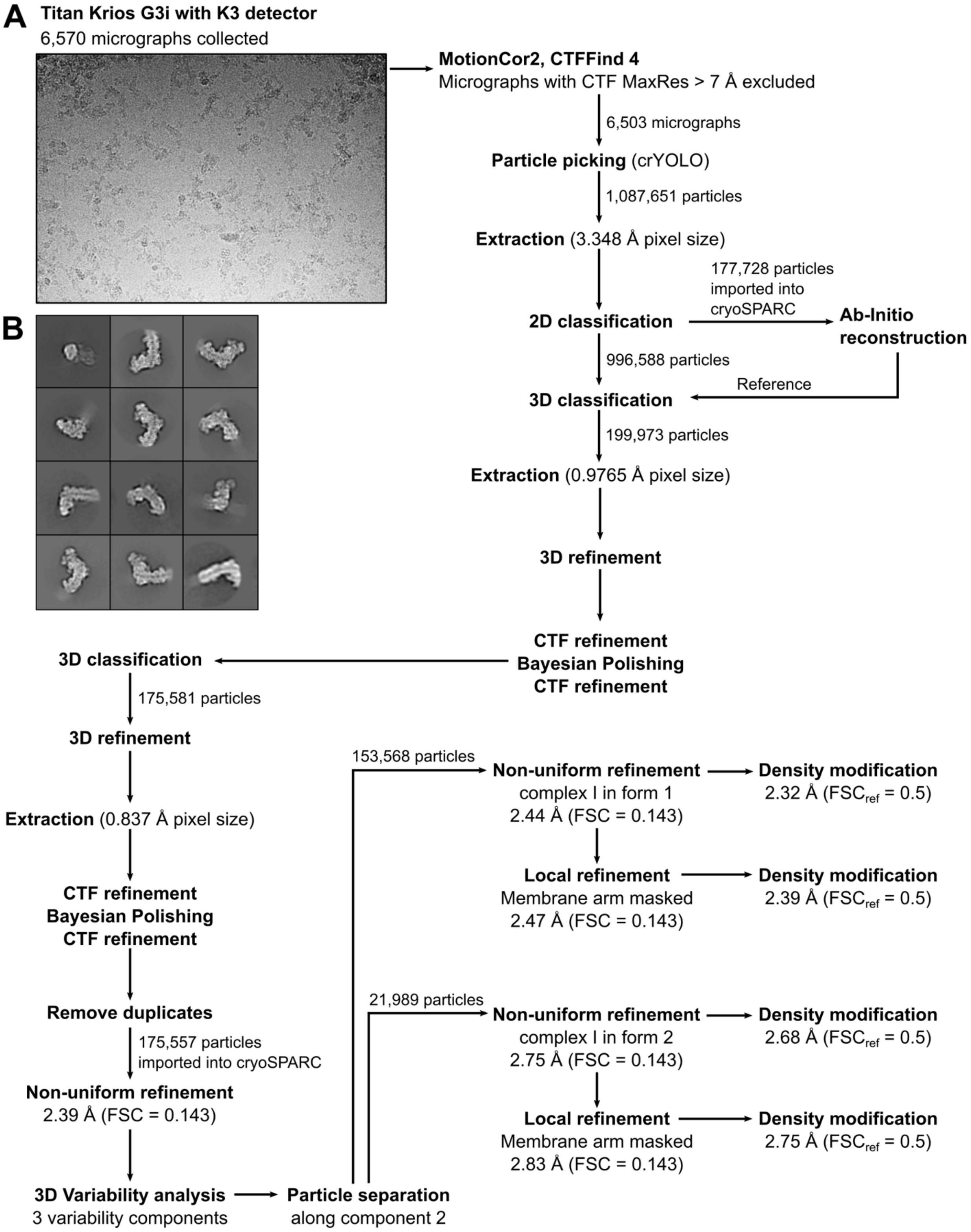


**Fig. S13. CryoEM processing pipeline for complex I in LMNG. (A)** Representative micrograph and workflow of single-particle cryoEM image processing of complex I solubilized in LMNG. **(B)** Representative 2D class averages.

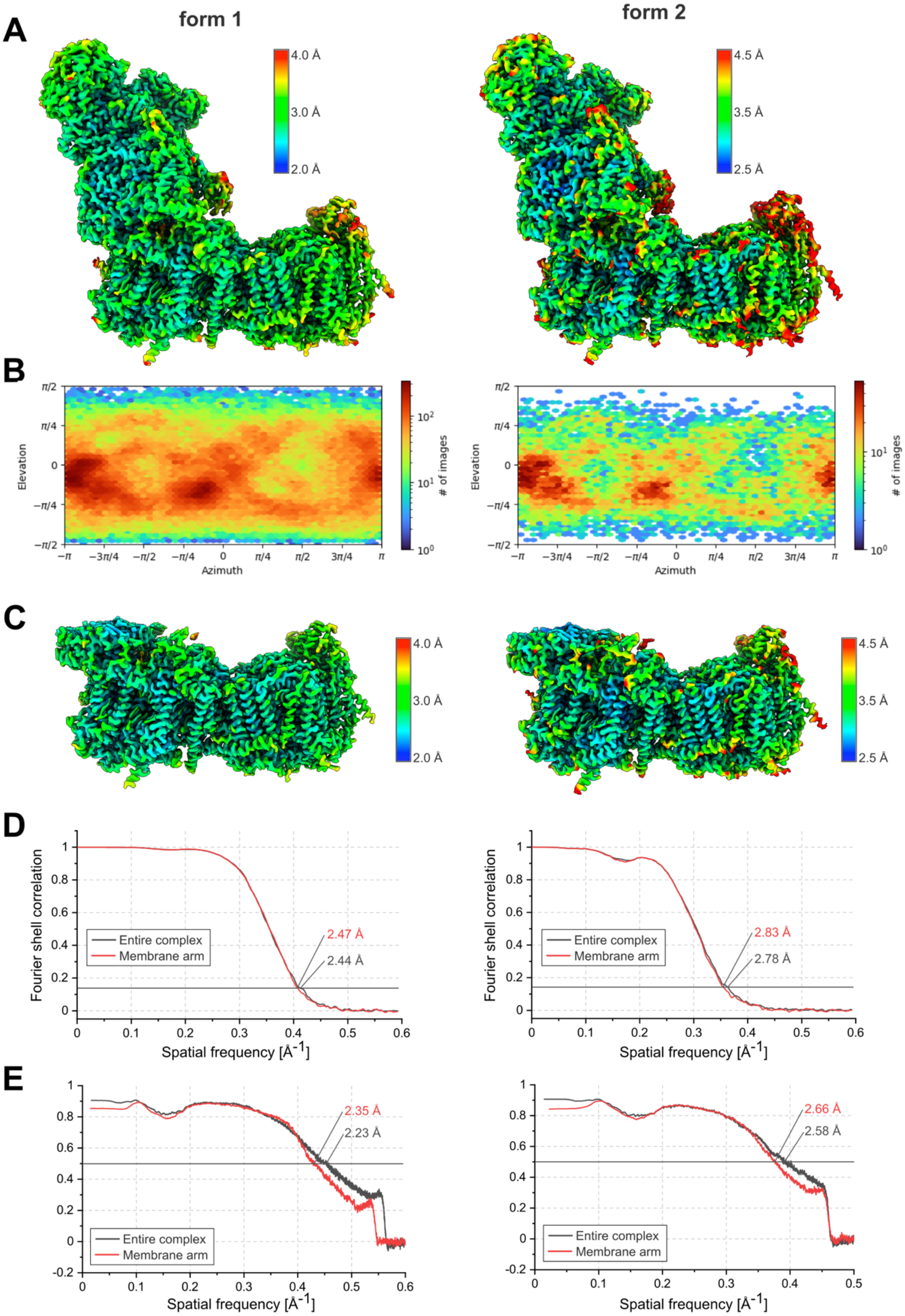
A
form 1
form 2
4.0 Å
3.0 Å
2.0 Å
4.5 Å
3.5 Å
2.5 Å
B
Elevation
Azimuth
### of images
C
D
Fourier shell correlation
Spatial frequency [Å-1]
Entire complex
Membrane arm
2.47 Å
2.44 Å
2.83 Å
2.78 Å
E
2.35 Å
2.23 Å
2.66 Å
2.58 Å

**Fig. S14. CryoEM processing of form 1 (left) and form 2 (right) of *Ct*- complex I in LMNG. (A)** Local resolution of entire map after non-uniform refinement. **(B)** Distribution of azimuth and elevation angles for particles included in the reconstruction. **(C)** Local resolution of the membrane arm of *Ct*-complex I after local refinement. **(D)** Two-half-map Fourier shell correlation curves. **(E)** Model-map Fourier shell correlation curves.

**Table S1. Subunits of *C. thermophilum* complex I**

| Subunit | Bovine nomenclature | Chain | Subunit | Bovine nomenclature | Chain |
|---|---|---|---|---|---|
| ***Membrane arm, core subunits*** | | | ***Peripheral arm, core subunits*** | | |
| ND1 | ND1 | 1 | NDUFS1 | 75-kDa | A |
| ND2 | ND2 | 2 | NDUFS2 | 49-kDa | C |
| ND3 | ND3 | 3 | NDUFS3 | 30-kDa | G |
| ND4 | ND4 | 4 | NDUFS7 | PSST | K |
| ND4L | ND4L | L | NDUFS8 | TYKY | I |
| ND5 | ND5 | 5 | NDUFV1 | 51-kDa | B |
| ND6 | ND6 | 6 | NDUFV2 | 24-kDa | H |
| ***Membrane arm, accessory subunits*** | | | ***Peripheral arm, accessory subunits*** | | |
| NDUFA1 [a] | MWFE | D | NDUFA2 | B8 | f |
| NDUFA3 | B9 | g | NDUFA5 | B13 | F |
| NDUFA8 | PGIV | U | NDUFA6 | B14 | P |
| NDUFA11 | B14.7 | J | NDUFA7 | B14.5a | Z |
| NDUFA13 | B16.6 | W | NDUFA9 | 39-kDa | E |
| NDUFB3 | B12 | c | NDUFA12 | B17.2 | h |
| NDUFB4 | B15 | j | NDUFAB1α | SDAP2 | O |
| NDUFB7 | B18 | 8 | NDUFS4 | 18-kDa | Y |
| NDUFB9 | B22 | R | NDUFS6 | 13-kDa | M |
| NDUFB11 | ESSS | S | *Ct*-NDUFV3 | 9-kDa | o |
| NDUFAB1β | SDAP1 | Q | | | |
| NDUFB2 | AGGG | e | | | |
| NDUFB5 | SGDH | n | | | |
| NDUFB6 | B17 | i | | | |
| NDUFB8 | ASHI | a | | | |
| NDUFB10 | PDSW | d | | | |
| NDUFC2 | B14.5b | b | | | |
| NDUFS5 | 15-kDa | 9 | | | |
| NUXM [b] | | X | | | |

[a] Last 5 residues of C-terminus modelled as poly-Ala

[b] Not present in mammalian complex I, *Y. lipolytica* nomenclature

**Table S2. Translation table for complex I residues from different species.**

Complex I subunit residues from *Chaetomium thermophilum* and corresponding residues in complex I from *Yarrowia lipolytica*, *Ovis aries*, *Homo sapiens* and *Thermus thermophilus*. Different subunit nomenclatures in *O. aries* and *T. thermophilus* are indicated in parentheses. *Ct*-complex I is compared with the ovine complex I structure (*12*). Residue positions in the published structure differ from the position in the subunit sequence in NDUFS2 and NDUFS7 as indicated in parentheses.

| **Subunit** | ***C. thermophilum*** | ***Y. lipolytica*** | ***O. aries*** | ***H. sapiens*** | ***T. thermophilus*** |
|---|---|---|---|---|---|
| ND1 (ND1, Nqo8) | Y151 | Y146 | Y142 | Y142 | Y162 |
| | E152 | E147 | E143 | E143 | E163 |
| | E201 | E196 | E192 | E192 | E213 |
| | E211 | E206 | E202 | E202 | E223 |
| | E236 | E236 | E227 | E227 | E248 |
| ND3 (ND3, Nqo7) | E39 | E39 | E38 | E38 | E45 |
| | C40 | C40 | E39 | E39 | S46 |
| | L64 | L64 | L63 | L63 | I69 |
| | D67 | D67 | D66 | D66 | D72 |
| ND4L (ND4L, Nqo11) | E30 | E30 | E34 | E34 | E32 |
| | E66 | E66 | E70 | E70 | E67 |
| ND6 (ND6, Nqo10) | E98 | E84 | E81 | E80 | E80 |
| | Y77 | Y63 | Y60 | Y59 | Y59 |
| | F85 | F71 | F68 | F67 | F67 |
| NDUFS2 (49-kDa, Nqo4) | H124 | H91 | H88 (H55) | H88 | H34 |
| | H128 | H95 | H92 (H58) | H92 | H38 |
| | Y177 | Y144 | Y141 (Y108) | Y141 | Y87 |
| NDUFS7 (PSST, Nqo6) | R128 | R108 | R114 (R77) | R111 | R69 |
| | R132 | R112 | R118 (R81) | R115 | R73 |

**Table S3. Data collection statistics**

| | CxI in LMNG (form 1) | CxI in LMNG (form 2) | CxI in DDM |
|---|---|---|---|
| **Data collection and processing** | | | |
| Microscope | Titan Krios G3i | Titan Krios G3i | Titan Krios G3i |
| Camera | K3 (counting) | K3 (counting) | K3 (counting) |
| Magnification | 105,000x | 105,000x | 105,000x |
| Calibrated pixel size [Å] | 0.837 | 0.837 | 0.837 |
| Voltage [kV] | 300 | 300 | 300 |
| Total exposure [$e^-/Å^2$] | 45 | 45 | 45 |
| Number of frames | 45 | 45 | 45 |
| Defocus range [µm] | -0.5 to -3.0 | -0.5 to -3.0 | -1.0 to -3.5 |
| Symmetry imposed | C1 | C1 | C1 |
| Number of micrographs | 6,503 | 6,503 | 1,996 |
| Initial particle images | 1,087,651 | 1,087,651 | 126,638 |
| Final particle images | 153,568 | 21,989 | 37,767 |
| Final resolution [Å] | | | |
| At FSC threshold 0.143, entire complex | 2.44 | 2.75 | 2.77 |
| At FSC threshold 0.143, membrane arm | 2.47 | 2.83 | 2.76 |
| Resolution after *DenseMod* [Å] | | | |
| At $FSC_{ref}$ threshold 0.5, entire complex | 2.32 | 2.68 | 2.75 |
| At $FSC_{ref}$ threshold 0.5, membrane arm | 2.39 | 2.75 | 2.75 |

**Table S4. Model statistics**

| | **CxI in LMNG (form 1)** | **CxI in LMNG (form 2)** | **CxI in DDM** | **CxI in LMNG (form 1, membrane arm)** | **CxI in LMNG (form 2, membrane arm)** | **CxI in DDM (membrane arm)** |
|---|---|---|---|---|---|---|
| Initial model used (PDB ID) | 6RFR, 6RFQ | 6RFR, 6RFQ | 6RFR, 6RFQ | 6RFR, 6RFQ | 6RFR, 6RFQ | 6RFR, 6RFQ |
| Refinement package | COOT, PHENIX real-space | COOT, PHENIX real-space | COOT, PHENIX real-space | COOT, PHENIX real-space | COOT, PHENIX real-space | COOT, PHENIX real-space |
| PDB accession | 7ZMG | 7ZMB | 7ZM7 | 7ZMH | 7ZME | 7ZM8 |
| Model resolution [Å] | | | | | | |
| At FSC threshold 0.5 | 2.23 | 2.58 | 2.69 | 2.35 | 2.66 | 2.68 |
| Model composition | | | | | | |
| Nonhydrogen atoms | 70,826 | 69,496 | 69,158 | 37,605 | 36,964 | 37,030 |
| Protein residues | 8,355 | 8,354 | 8,389 | 4,390 | 4,396 | 4,429 |
| Ligands | 1 Zn<br>6 SF4<br>2 FES<br>1 FMN<br>1 NDP<br>2 ZMP<br>5 CDL<br>9 PC1<br>18 3PE<br>3 LMN | 1 Zn<br>6 SF4<br>2 FES<br>1 FMN<br>1 NDP<br>2 ZMP<br>5 CDL<br>8 PC1<br>17 3PE<br>3 LMN | 1 Zn<br>6 SF4<br>2 FES<br>1 FMN<br>1 NDP<br>2 ZMP<br>5 CDL<br>4 PC1<br>8 3PE<br>14 LMT | 1 ZMP<br>5 CDL<br>8 PC1<br>17 3PE<br>3 LMN | 1 ZMP<br>5 CDL<br>7 PC1<br>16 3PE<br>3 LMN | 1 ZMP<br>5 CDL<br>4 PC1<br>7 3PE<br>14 LMT |
| Waters | 2,649 | 1,593 | 1,219 | 1,111 | 633 | 608 |
| Cross-correlation | | | | | | |
| Mask | 0.81 | 0.81 | 0.81 | 0.86 | 0.85 | 0.83 |
| Volume | 0.77 | 0.76 | 0.78 | 0.80 | 0.79 | 0.78 |
| B-factors [Å$^2$] (mean) | | | | | | |
| Protein | 27.13 | 17.34 | 22.72 | 18.85 | 27.54 | 12.88 |
| Ligand | 38.17 | 26.47 | 30.29 | 25.38 | 35.34 | 22.82 |
| Water | 11.10 | 8.94 | 13.55 | 9.96 | 16.32 | 12.65 |
| R.m.s deviations | | | | | | |
| Bond lengths [Å] | 0.008 | 0.009 | 0.012 | 0.008 | 0.010 | 0.010 |
| Bond angles [°] | 1.277 | 1.342 | 1.462 | 1.206 | 1.417 | 1.509 |
| Validation | | | | | | |
| MolProbity Score | 1.85 | 1.85 | 1.78 | 1.84 | 1.66 | 1.65 |
| Clash score | 9.05 | 11.34 | 9.92 | 9.65 | 8.85 | 8.06 |
| Poor rotamers [%] | 1.81 | 0.84 | 0.85 | 1.84 | 0.67 | 0.72 |
| Ramachandran plot | | | | | | |
| Favored [%] | 97.02 | 95.92 | 96.19 | 97.25 | 96.89 | 96.66 |
| Allowed [%] | 2.97 | 4.05 | 3.79 | 2.70 | 3.11 | 3.27 |
| Disallowed [%] | 0.01 | 0.04 | 0.02 | 0.05 | 0.00 | 0.07 |

**Movie S1**
Reconstruction series from 3D variability analysis of DDM-solubilized *Ct*-complex I

**Movie S2**
Reconstruction series from 3D variability analysis of LMNG-solubilized *Ct*-complex I.

**Movie S3**
Reconstruction series from 3D variability analysis of LMNG-solubilized *Ct*-complex I with mask application at hinge region.

**Movie S4**
Atomic model morph of *Ct*-complex I from form 1 to form 2 at the π-gate.
